# Structural Plasticity of RRE Stem-Loop II Modulates Nuclear Export of HIV-1 RNA

**DOI:** 10.1101/2025.02.20.639096

**Authors:** Manju Ojha, Lucia Hudson, Amanda Photenhauer, Trinity Zang, Lauren Lerew, Şölen Ekesan, Jason Daniels, Megan Nguyen, Hardik Paudyal, Darrin M. York, Melanie D. Ohi, Jan Marchant, Paul D. Bieniasz, Deepak Koirala

**Affiliations:** Department of Chemistry and Biochemistry, University of Maryland Baltimore County; Baltimore, MD, USA; Life Sciences Institute, University of Michigan; Ann Arbor, MI, USA; Laboratory of Retrovirology, The Rockefeller University, New York, NY, USA; Department of Chemistry and Chemical Biology, Laboratory for Biomolecular Simulation Research, and Institute for Quantitative Biomedicine, Rutgers University; Piscataway, NJ, USA; Department of Cell and Developmental Biology, University of Michigan; Ann Arbor, MI, USA

## Abstract

The Rev Response Element (RRE) forms an oligomeric complex with the viral protein Rev to facilitate the nuclear export of intron-retaining viral RNAs during the late phase of HIV-1 infection. However, our structural understanding of this crucial virological process remains limited. In this study, we determined several crystal structures of an intact RRE stem-loop II in two distinct conformations, performed negative-staining electron microscopy and molecular dynamics simulations, and revealed that this three-way junction RNA exhibits remarkable structural plasticity. Through *in vitro* Rev-binding and *in vivo* Rev-activity assays using various stem-loop II mutants designed to favor one of the conformers, we demonstrated that the structural plasticity of stem-loop II modulates Rev binding and oligomerization. Our findings illuminate emerging perspectives on RRE dynamics-based regulation of HIV-1 RNA nuclear export and provide a framework for developing anti-HIV drugs that target specific RRE conformations.

## MAIN

The human immunodeficiency virus type 1 (HIV-1) consists of about 9 kb long genomic RNA that encodes the proteins required for the virus proliferation in the host cells ^1–3^. During infection, the proviral DNA integrated within the host genome produces a transcript that undergoes alternative splicing or remains unspliced to yield intron-less or intron-retaining RNAs. These RNAs are eventually exported to the cytoplasm to be translated into viral proteins or packaged into new virions ^1–3^. Because the host nuclear export machinery does not transport unspliced or intron-retaining RNAs, several retroviruses, including HIV-1, have evolved specific mechanisms to promote the nuclear export of such RNA species ^4–6^. In HIV-1, a viral protein called Rev recognizes a unique *cis-acting* RNA structure called the Rev Response Element (RRE) found within the intronic regions of the unspliced or partially spliced RNAs ^5–9^. Such RRE-Rev assembly then interacts with the host machinery to guide the nuclear export of these viral RNAs through the Crm1 pathway ^10–12^. As this RRE-Rev platform is essential for viral proliferation, it has been an attractive target for therapeutic interventions against HIV-1 infection ^13–15^. However, the molecular mechanisms of RRE-mediated nuclear export of HIV-1 RNAs remain elusive, mainly due to the lack of high-resolution structures of the RRE and well-defined structural models for its interactions with Rev. Here, we determined crystal structures of the high-affinity Rev binding site of the HIV-1 RRE, the intact stem-loop II (SLII), in two alternative conformations using Fab-assisted crystallography. Through *in vitro* Rev-binding assays, structure-guided HIV-1 nuclear export activity measurements in human cells, electron-microscopy, and computational dynamics simulation approaches, we demonstrate that the structural plasticity captured in the crystal structures is vital for regulating HIV-1 Rev oligomerization and nuclear export.

HIV-1 RRE comprises ∼350 nucleotides ^2, 7–9^. However, previous computational, enzymatic, and chemical probing experiments with mutant and truncated constructs have identified that ∼232 nucleotides are sufficient to form the functional core of the RRE ^16, 17^. The proposed secondary structure comprises five subdomains: I to V, with II to V being the stem-loops (SLs) that branch out from base stem I (Figure 1A), and SLII is known to form a three-way junction (3WJ) structure, where the base stem IIa bifurcates into stem loops IIb and IIc. Within this RRE secondary structural model, an asymmetric purine-rich bulge within the stem I and the 3WJ region of SLII have been identified as the high-affinity binding sites for Rev ^18–24^. While it is known that multiple Revs bind to the RRE to form a homo-oligomeric ribonucleoprotein (RNP) complex, how different structural features within the RRE orchestrate cooperative binding of Rev oligomers to construct a functional RRE-Rev complex remains unclear. For instance, significant alterations, such as the complete deletion of SLII, eliminate Rev binding and nuclear export activity, whereas disruption of the stem-loops III and IV do not exhibit such a drastic effect ^25–28^.

**Figure 1.**
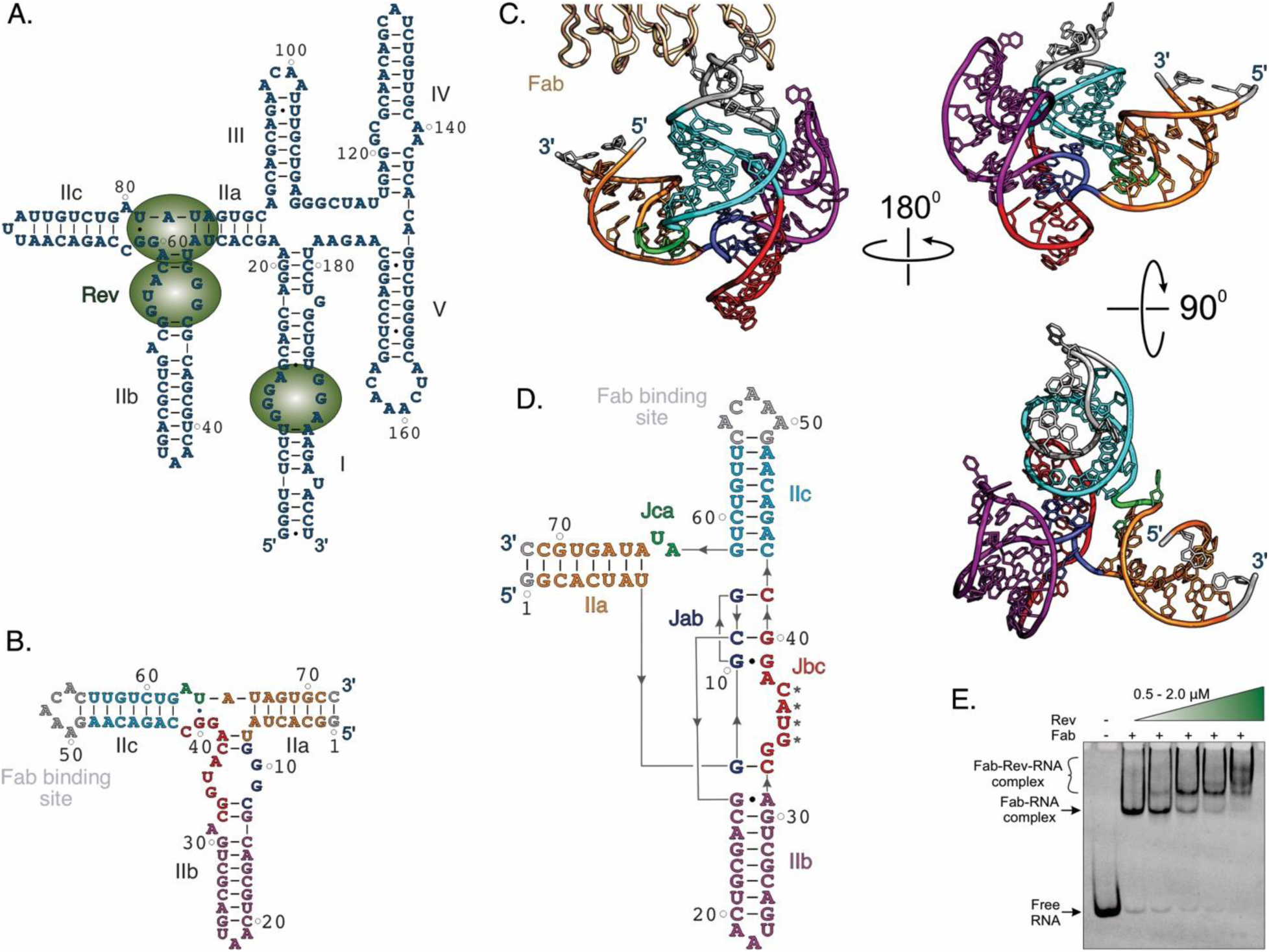
The overall structure of HIV-1 RRE SLII crystallized with a Fab chaperone. (A) A secondary structural model of a minimal HIV-1 RRE with five stem-loop structures. The green spheres represent the Rev molecules at the previously identified high-affinity binding sites. (B) A secondary structure model of the SLII crystallization construct (SLIIc) in which the Fab BL3-6 binding epitope GAAACAC sequence replaces the IIc loop. The gray nucleotides indicate the mutations or insertions compared to the SLII wild-type sequence. (C) The crystal structure of the SLIIc cocrystallized with the Fab and determined at 2.42 Å resolution and (D) its crystal-derived secondary structure. For clarity, the Fab is obscured in the rotated views. The asterisks in panel D depict the nucleotides involved in crystal contacts. (E) A native polyacrylamide gel showing the binding of the Fab with the crystallization construct without perturbing the Rev binding. The green color-gradient triangle shows the Rev concentration ranging from 0.5 to 2.0 µM with the 0.5 µM increment in each lane. All figure panels and corresponding labels, if any, are colored analogously for facile comparison.

Several studies have investigated the structures and interactions of HIV-1 RRE to illuminate the possible mechanism of RRE structure-mediated nuclear export of HIV-1 RNA species. Phylogenetic and biochemical probing studies have provided secondary structure information on HIV-1 RRE, including base pairings and nucleotide accessibility ^20, 24, 29–31^. Results from these studies led to a model where the full-length HIV-1 RRE can adopt multiple conformations, with the five and four stem-loop conformations being the most prevalent forms ^24, 31, 32^. Conformational diversity has been shown to originate due to the structural rearrangements of the stem-loops III and IV; however, SLII, the high-affinity Rev binding site, remained virtually unchanged in those models ^2, 24, 30–32^. Therefore, SLII has been the focus of several biochemical and structural studies to understand the nature of RRE-Rev interactions and the RRE-mediated HIV-RNA nuclear export mechanism. NMR and X-ray crystallographic studies of isolated IIb constructs alone and in complex with a Rev peptide not only illuminated how Rev binds to the RNA through base-specific hydrogen bonds and electrostatic interactions with the phosphodiester backbone but also underscored that this RNA likely undergoes some conformational changes upon Rev binding ^33–38^, indicating a structurally plastic nature of the RNA. Such plasticity at the SLII 3WJ seemed necessary for the cooperative binding of multiple Rev molecules ^21–23, 35, 38^, a model further supported by a crystal structure of the isolated IIb construct with an engineered junction site in a complex with the Rev dimer ^37^. It is also consistent with high SHAPE reactivities observed for the IIb near the 3WJ nucleotides in the context of the intact RRE sequence, which agrees with the highly flexible nature of the Rev binding sites ^2, 31, 32^. These observations with the isolated IIb structures indicate the possibility of similar conformational flexibility for the intact SLII 3WJ, possibly modulating the cooperative Rev binding to the HIV-1 RRE. However, other than structures of select SLII fragments ^33–38^, a SAXS-based study proposing a low-resolution model with an unusual A-shaped topology for an intact HIV-1 RRE ^39^ and a recently reported SLII crystal structure ^40^, the nature of RRE structural plasticity and how it modulates Rev interactions and mechanism of HIV-1 RNA nuclear export remains largely unknown.

Here, we report multiple high-resolution crystal structures of intact SLII constructs in two alternative forms: compact and extended conformations. We reveal that these SLII conformers fold into unique architectures with a distinct organization of the 3WJ. One of the SLII conformers exhibits extensive interactions between the 3WJ junction-forming nucleotides with a tight arrangement of the stems around the 3WJ, designated as the compact form. The second SLII conformer folds into a more open arrangement and elongated IIb, termed the extended form. Although both conformers may coexist in solution, our findings from combined negative-stain electron microscopy (EM), molecular dynamics analysis, and biochemical Rev-binding assays show that, while the extended conformation seems thermodynamically favored in solution, the compact conformation of SLII, stabilized by engineered mutations, supports more robust Rev binding and oligomerization compared to the extended form. Furthermore, the compact form is consistent with a Rev-bound conformation *in vivo* that promotes nuclear export, as indicated by Rev activity measurements in HIV-1 transfected human cells. Thus, our findings suggest a model where the plasticity of the SLII could act as a structural regulator to modulate Rev binding cooperativity and oligomerization to the HIV-1 RRE, shedding light on the Rev-RRE-mediated HIV-1 nuclear RNA export mechanism and providing unique opportunities to develop RRE conformation-specific anti-HIV therapeutics.

## RESULTS

### SLII crystals with a Fab chaperone diffracted to 2.42 Å resolution

Our crystallization constructs span 68 nucleotides (A22 to A89) of the HIV-1 RRE that cover the intact 3WJ region of the SLII (Figure 1A). These constructs include A22G and A89C mutations that install a G-C pair to close the IIa stem for efficient transcription and stability. Crystallization of the wild-type construct (SLII_WT_) was unsuccessful, so we turned our efforts to Fab-assisted RNA crystallography ^41–44^. We prepared two crystallization constructs, SLIIb (Figure S1) and SLIIc (Figure 1B), by replacing the IIb and IIc loops with Fab BL3-6 binding hairpin-loop sequence 5′ gAAACAc. The native polyacrylamide gel electrophoresis (nPAGE) assay showed that a recombinantly expressed Rev (see Methods) binds to both crystallization constructs similarly compared to the SLII_WT_ construct (Figure 3 below and S2), suggesting that the engineered Fab-binding sequence did not influence the overall folding of the SLII RNA. More importantly, the Rev also binds to these constructs in the presence of the Fab BL3-6, forming a ternary complex (Figures 1E, S2), supporting that the Fab binding site in these constructs is away from the Rev binding site. Thus, the Fab binding is less likely to alter the overall folding of the SLII crystallization constructs. Following the Fab and Rev binding tests, we set up the crystallization trials of both SLIIb and SLIIc constructs in complex with the Fab. However, only the SLIIc-Fab complex yielded robust crystals that diffracted to 2.42 Å resolution. The crystallographic data suggested that the SLIIc-Fab complex was crystallized in the P1 space group and contained two complexes within the crystallographic asymmetric unit (Table S1 details the data collection statistics). The initial phases for solving the structure were obtained via molecular replacement using a previous crystal structure of Fab BL3-6 (PDB: 8T29) ^43^ as a search model that provided a robust electron density map, allowing the modeling of the SLII nucleotides unambiguously. After iterative rounds of model building and refinements, the final structure was determined at 2.42 Å resolution, which converged with *R_work_* and *R_free_* values of 20.5% and 26.5%, respectively (Table S1 details of the refinement statistics). The two SLIIc RNA copies within the crystallographic asymmetric unit appear very similar (all atoms alignment root mean square deviation, RMSD = 2.55 Å) except for some deviations in the relative orientation of the IIb helix (Figure S3), perhaps due to the high flexibility and dynamicity of this region, which is consistent with relatively weaker electron density map and high crystallographic B-factors observed for this region (Figure S4).

**Figure 2.**
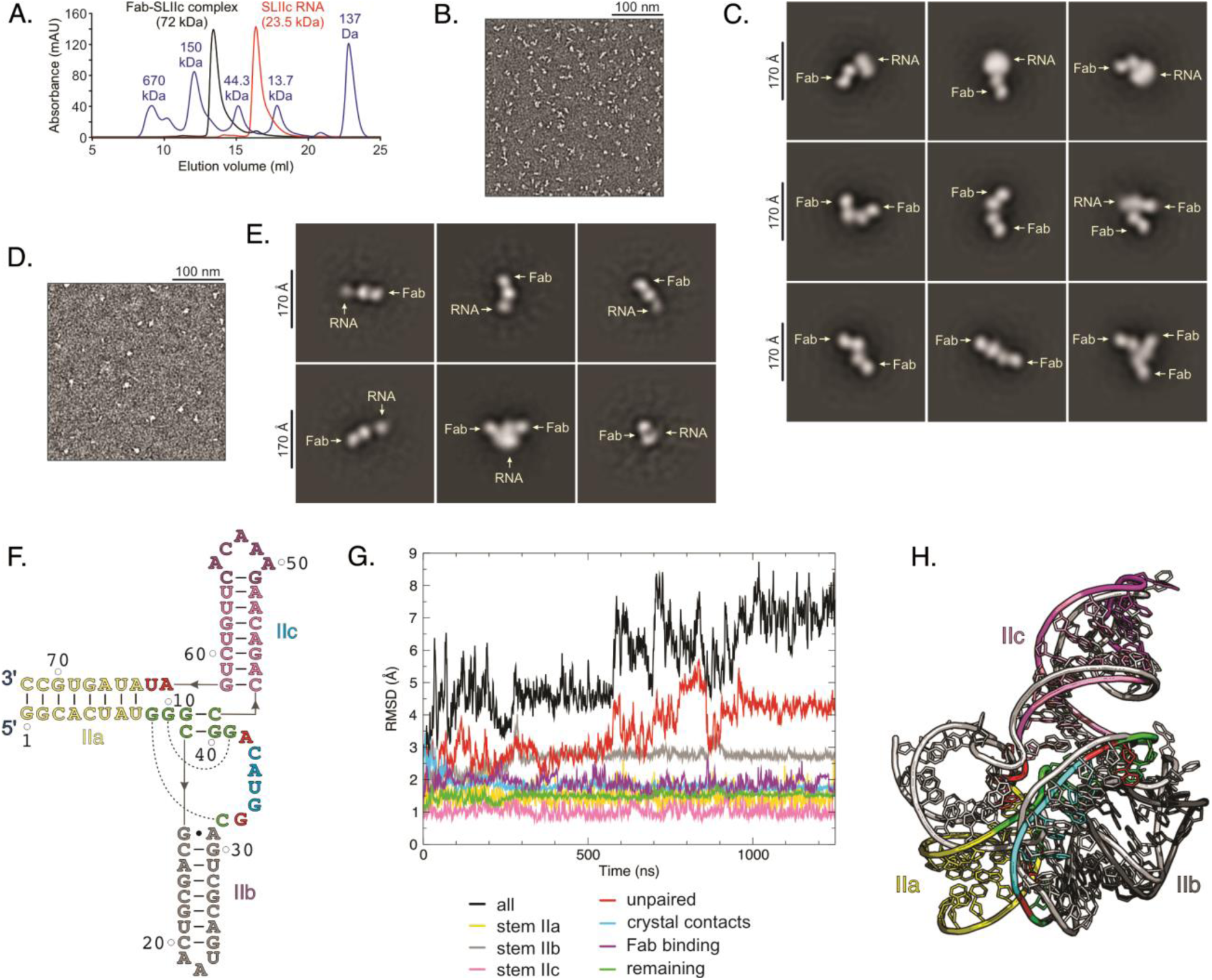
Size-exclusion chromatography, negative-staining EM and MD simulation studies of the HIV-1 RRE SLII. (A) Representative chromatograms for the SLIIc (red), Fab-SLIIc complex (black) and a molecular size marker (blue) suggest no dimerization of the SLIIc or its Fab complex. (B) Representative negatively stained image of the Fab-SLIIc complex particles at a 1.2 µg/ml concentration. (C) Representative 2D averages for this concentrated sample show monomers, dimers, and trimers of the Fab-SLIIc complex. The oligomerization of the complexes at this concentration is not homogenous. (D) Representative negatively stained image of the same sample at a 0.24 µg/ml concentration. (E) Representative 2D averages showing the Fab-SLIIc monomers in the dilute solution. The sample at a more dilute concentration is predominantly monomeric. (F) The crystal-derived secondary structure for the MD simulation. The stems IIa-c are colored yellow, gray, and pink, respectively. The unpaired nucleotides are colored red, and the nucleotides taking part in crystal contact are colored cyan. The nucleotides that are sequentially not part of the stem but structurally stacked on the stems are colored green, and the nucleotides involved in the Fab-binding are colored purple. The dotted curves represent long-range base-pairing interactions. (G) The root mean square deviations (RMSDs) of different structural components compared to the starting structure. Along the 1.25 µs (1250 ns) trajectory, frames were collected every 10 picoseconds, and RMSDs were plotted as running averages over 100 frames (1 ns). (H) The superposition of the final simulated structure (colored according to the secondary structure as shown in F) with the crystal structure (gray).

**Figure 3.**
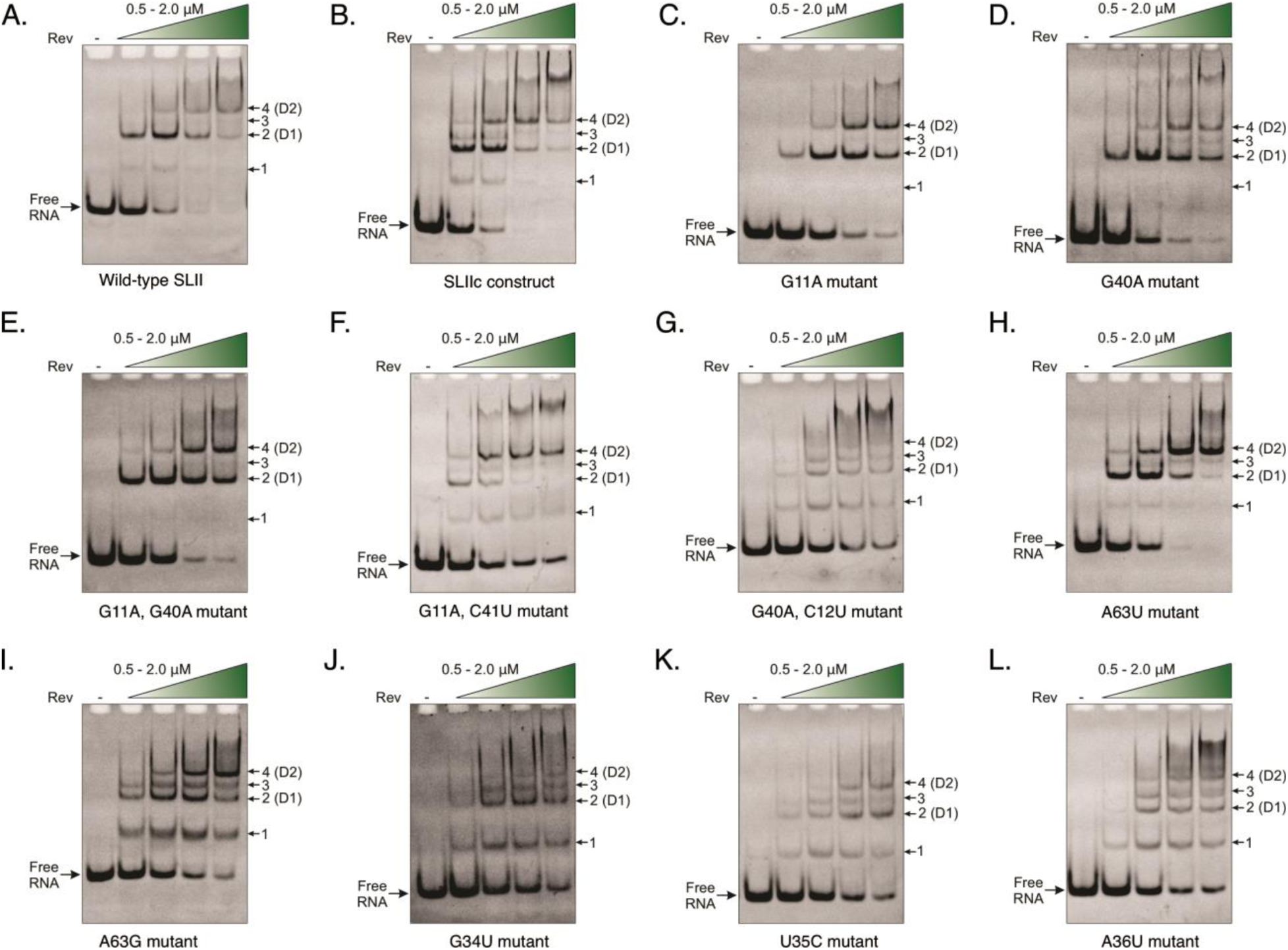
Binding cooperativity and oligomerization of the HIV-1 Rev within the SLII 3WJ. Native polyacrylamide gels showing the Rev-binding and oligomerization with the (A) wild-type SLII and (B) crystallization construct (SLIIc). Similar native gels for Rev-binding with (C) G11A; (D) G40A; (E) G11A, G40A; (F) G11A, C41U; (G) G40A, C12U; (H) A63U; (I) A63G; (J) G34U; (K) U35C; and (L) A36U mutant SLIIc constructs. The green color-gradient triangle shows Rev concentration ranging from 0.5 to 2.0 µM with the 0.5 µM increment in each lane.

### HIV-1 RRE SLII adopts an intricate 3WJ fold

The SLII crystal structure adopted roughly a Y-shaped topology, in which the base stem IIa bifurcates into the IIb and IIc stems to organize a 3WJ fold (Figures 1C, S3, S4). The IIa stem resides almost perpendicular to the IIc stem, and IIb protrudes from the IIa-IIc junction, leaning towards the IIc with the IIb loop near the 3WJ region. As shown in the crystal-derived secondary structure (Figure 1D), starting from the 5′ end, the nucleotides G1-U8 base pair with C72-A65 to form the IIa stem, and G9-C12 compose the junction (Jab) between IIa and IIb. The base pairing of G13-C20 with G24-A31, including a non-canonical G13•A31 pair bordering the 3WJ, forms the stem IIb, which is capped by the trinucleotide A21A22U23 loop. The sequence C32-C41 comprises the IIb-IIc junction (Jbc) interacting with the Jab nucleotides via canonical and noncanonical base pairing. Following the Jbc, the C42-G49 base pairs with the C55-G62 to form the IIc stem, which is capped by the Fab binding loop used for the crystallization. Finally, the junction Jca connects the IIc and IIa helices and contains two unpaired nucleotides (A63 and U64).

The 3WJ in SLIIc crystal structure appeared complicated but well-structured, with extensive interactions between the junction nucleotides, suggesting a prominent role of this junction for Rev interactions (see Figure S5 for details). The four nucleotides within the Jab form a sharp turn in the helix, allowing the G9 to base-pair with the Jbc C32 and G10, G11, and C12 to base-pair with the Jbc G39, C41 and G40, respectively. Including the G10•G39 non-canonical pair, the three base-paired helix remains co-axially stacked with the IIc, forming almost a continuous A-form helical stem. Whereas the A38 and G39 and G33 and C32 remain helically stacked near the IIb and IIc stems, respectively, the nucleotides G34 to C37 flip out and form base pairs with the palindromic sequence of the symmetry-related molecule (Figure S6). Besides stacking interactions, the unpaired nucleotides G33 and A38 are also stabilized by hydrogen bonding interactions. The 3WJ junction connects with the stem IIb via the G13•A31 non-canonical pair, which stacks co-axially on the G9 and C32 pair. Within the Jca, the A63 and U64 nucleotides are unpaired, where U64 flips out, and A63 is stabilized through stacking interactions with the U8 and A65 base pair within the IIa stem. While the Jca nucleotides are not engaged in interactions with other junction nucleotides, the asymmetric number of the junction nucleotides (Jbc > Jab > Jca) and extensive interactions between the Jab and Jbc nucleotides define the relative positioning of the helices flanking the 3WJ. Overall, unlike a relatively simple 3WJ with several non-canonical pairs as predicted based on the NMR and crystal structures of the IIb constructs ^37, 38^, the 3WJ in our crystal structure is stabilized by networks of base-stacking and hydrogen bonding interactions with canonical base pairs, resulting in a more extensive and well-organized junction in the context of the full-length SLII. However, these unpaired nucleotides underscore the high dynamicity and propensity of the 3WJ to adopt multiple conformations.

### EM and MD simulations provide evidence of SLII crystal structure in solution

The structural features around the 3WJ do not involve any interactions with the Fab in the crystal lattice, and Fab did not inhibit

Rev binding (Figure 1E), indicating that the overall folding of our crystal structure is unlikely to be affected by complex formation with the Fab. Additionally, while the tertiary structure of the 3WJ fold appears complicated, the base-paired nucleotides of the IIa, IIb and IIc helices and the formation of IIb and IIc loops are consistent with previous biochemical, enzymatic probing and SHAPE-based studies (see Figure S7) in the context of the full-length HIV-1 RRE or the complete HIV-1 genome ^2, 31, 32, 45^. Nevertheless, the four nucleotides, G34-C37, which comprises the putative Rev binding site, were observed to form base-pairing crystal contacts with the identical nucleotides of the symmetry-related SLIIc molecules (Figure S6). Therefore, we performed several experiments to determine whether these interactions exist in solution. First, SEC analyses of the SLIIc and Fab-SLIIc complex showed single homogeneous peaks within the expected monomeric sizes compared to a standard marker, indicating that the crystalized construct does not dimerize in solution (Figure 2A). Second, we performed negative-stain EM analysis with concentrated and dilute Fab-RNA samples. While the classification of particles isolated from images of negatively stained particles in concentrated solution showed 2D classes of what appear to be the Fab-RNA complexes as monomers, heterogeneous dimers, and trimers (Figures 2B, C, and S8), the 2D averages generated from particles isolated from images of negatively stained particles in the more dilute sample showed that almost all the classes were single Fab-RNA complexes, with only a few classes that appeared to be dimerized complexes (Figures 2D, E and S9), showing that although Fab-RNA complexes can form heterogenous dimers and trimers in concentrated solutions, they exist as monomers in dilute solutions. Although low resolution, none of these classes match the topology of the dimer observed in the crystal lattice (Figure S6), further suggesting that the base-pairing of the G34-C37 nucleotides represents crystallographic contacts, and these interactions are likely irrelevant in solution. Third, we performed molecular dynamics (MD) simulations using AMBER22 ^46^. We examined the dynamical behavior of the nucleotides in crystal contacts compared to those in the stems using MD simulations departing from the crystal structure and monitoring the RMSDs (Figures 2F-G, S10, S11). The RMSD analyses indicated significant mobility of the relative orientations of stems IIa, IIb, and IIc rather than fluctuations occurring internally to these stems (Figures 2G, S10). While crystal contact forming nucleotides (G34-C37) displayed converged RMSD values overlapping in magnitude with that of the fully base-paired stem IIa, other unpaired nucleotides (G33, A38, A63, U64) exhibited high RMSD fluctuations correlating with the overall RMSD values, suggesting that internal structure of the stems, crystal contacts, and the Fab binding site are stable and remain close to the crystal structure during simulations, and thus the crystal structure represents at least a metastable free energy basin in solution (Figures 2G, H). Notably, the four crystal-contact-forming nucleotides in the simulated structure turned towards the RNA and formed non-canonical hydrogen bonds with nearby nucleotides – specifically, C37:O2 with A38:N6, A36:N6 with O2′ of G9 and C14, G34:N7 with G9:N2 (Figure S10). Surprisingly, the RMSD fluctuations for the G34-C37 and unpaired nucleotide group (G33, A38, G63, U64) exhibited different behaviors in the A63G mutant SLII simulations. Although this nucleotide is located far from the crystal contacts, this behavior is likely due to sampling slightly different basins or local rearrangements in the base-pairing patterns resulting from the mutation (Figures S10, S11).

### 3WJ comprises Rev binding and oligomerization sites

To understand the nature of SLII-Rev interactions, we performed binding studies with the SLIIc and mutant SLIIc constructs using nPAGE assays. We used a Rev construct with L12S and L60R mutations, which were shown to limit the Rev protein to dimerization ^47^. We also introduced two additional mutations, P28A and V16D, to this construct and included an N-terminal MBP1 tag to enhance Rev protein solubility. First, the binding assays suggested that SLII (SLII_WT_ and SLIIc) binds at least four copies of the Rev molecules, as indicated by four distinct Rev-SLII complex bands, and the nature of these bands implied that SLII has two different sites for Rev dimerization (Figures 3A, B). Whereas the Rev binding to the comparatively higher-affinity site at lower Rev concentrations likely formed a homodimeric complex with SLII (D1), the higher Rev concentrations enabled the second Rev homodimer to bind the relatively lower-affinity site (D2). Next, we introduced a series of rationally designed, base-paring disruptive or compensatory mutations within the 3WJ based on the SLIIc crystal structure and performed their Rev binding tests. Whereas G11A or G40A mutations expected to disrupt the central G40:C12 or C41:G11 pairs had a minimal effect on overall binding affinity (based on the band intensities for free RNA), the cooperativity between two Rev dimerization sites seemed to be more compromised as the Rev dimerization was more limited presumably to the D1 site even at higher Rev concentrations (Figures 3C, D). The G11A and G40A double mutation exhibited similar Rev binding patterns (Figure 3E), indicating that these nucleotides might have roles in regulating the cooperativity of Rev binding. Nonetheless, it is possible that these mutations disrupt the D1 site, and the complex formed by Rev binding at the D2 site migrates in the gel similarly to a complex formed at the D1 site. Consistently, the base pair compensatory mutations G11A and C41U to replace the G11:C41 by the A11:U41 pair rescued the binding cooperativity to a similar level to that of non-mutant SLIIc (Figure 3F), suggesting that this 3WJ stabilizing base-pair is important for high-affinity and cooperative Rev binding and oligomerization. Surprisingly, the compensatory G40A and C12U mutations that substitute the G40:C12 pairs with A40:U12 reduced the specific binding and dimerization of the Rev at both D1 and D2 sites (Figure 3G). While these nucleotides play roles in defining the specificity of Rev binding, such unprecedented results for these rationally designed mutants indicate potential rearrangements of interactions among the 3WJ nucleotides that retain a conformation pertinent to cooperative Rev binding and oligomerization, consistent with the highly plastic nature of this RNA. Although these mutational effects on reduced binding affinity and cooperativity are generally consistent with reduced Rev activity in previous *in vivo* measurements based on the translational repression assays ^48^, those assays did not explain the possible influence of SLII structural rearrangements on the observed Rev activities.

On the other hand, the A63U (Figure 3H) and A63G (Figure 3I) mutants within Jca showed similar Rev binding patterns as the non-mutant SLIIc (Figure 3B), suggesting that Jca is not the primary site for Rev binding and oligomerization. Notably, Jca nucleotides have no significant role in the 3WJ stabilization in the SLIIc crystal structure as they bulge out without any specific interaction with the junction nucleotides (Figures 1C, S5), which is consistent with no effect of this bulge on Rev activity observed in previous *in vivo* studies ^25, 49^. As primarily Rev binding sites are located within the Jab and Jbc, next, we mutated nucleotides observed in the crystal contacts and performed Rev-binding assays. The G34U, U35C and A36U mutations all showed reduced overall binding with Rev, and complex formation was somewhat more restricted to Rev dimerization at the D1 site even at higher Rev concentrations (Figures 3J, K and L, respectively), suggesting that these nucleotides might specifically interact with Rev and modulate the cooperativity of Rev oligomerization. While the reduced binding with these mutations correlates well with nominal Rev activities observed in previous *in vivo* studies ^48^, given several unpaired nucleotides around the SLIIc 3WJ, a single-nucleotide mutation is less likely to account for the loss of such a robust specificity and cooperativity of Rev binding and oligomerization.

### SLIIG34U mutant crystal structure adopts an alternate fold

Our binding studies with SLII mutants, especially the nucleotides involved in crystal contacts, suggest their crucial roles in defining the overall conformational state of the SLII that modulates Rev binding and oligomerization. To test this hypothesis, we crystallized and solved the structure of the G34U mutant at 3.0 Å resolution using the same Fab-assisted crystallography approach (see Figure S12 and Table S1 for statistics). Unlike the SLIIc-Fab complex, the crystallographic asymmetric unit contained a single Fab-RNA complex without significant RNA-RNA crystal contacts (Figure S13). Excitingly, the SLIIc G34U crystal structure appeared in a distinctly different architecture than the SLIIc structure, especially around the 3WJ, revealing substantial structural changes induced by this single G34U mutation (Figure 4A). In this mutant structure, the IIa stem resides nearly perpendicular to the IIc stem, and the IIb stem protrudes out from the IIa-IIc junction almost at the right angle (Figures 4A, B). The crystal-derived secondary structure showed that the organization of the 3WJ is relatively simple compared to its non-mutant SLIIc structure (Figure 4C). While the paired and unpaired nucleotides within the IIa and IIc stems along with the Jca are similar in SLIIc and SLIIc G34U structures, the reorganization of the base-pairing interactions between the Jab and Jbc nucleotides within the SLIIcG34U 3WJ form a more extended IIb stem than the SLIIc structure. Notably, the overall architecture of this extended IIb appears similar to the previously reported crystal structure of an isolated IIb construct ^37^. Within the SLIIcG34U 3WJ structure, Jab (G9-C12) has no unpaired nucleotides due to the G9:C37, G10•U35, G11•U34 and C12:G33 base pairs (Figure S14). Unlike the G13•A31non-canonical base pair in the SLIIc structure, this mutant structure formed the G13:C32 base pair, leaving the A31 unpaired and flipped out of the helix. Three of the four unpaired nucleotides G34-C37 in the SLIIc 3WJ are stabilized by the base-pairing in the SLIIcG34U 3WJ with the formation of a new unpaired nucleotides A38-C41 (Figures 4C, S14). It implies that Jbc G34-C37 nucleotides play crucial roles in defining the conformational state of the SLII, and the loss of Rev binding and oligomerization by the G34U mutation is most likely due to the re-registration of the G34-C37 interactions. The G11•U34 pair in the mutant structure would have been a G11•G34 non-canonical pair in its non-mutant form. This change destabilizes the extended 3WJ to switch to the more compact form of the SLIIc, as a G11•G34 is expected to be thermodynamically less stable than a G11•U34 pair.

**Figure 4.**
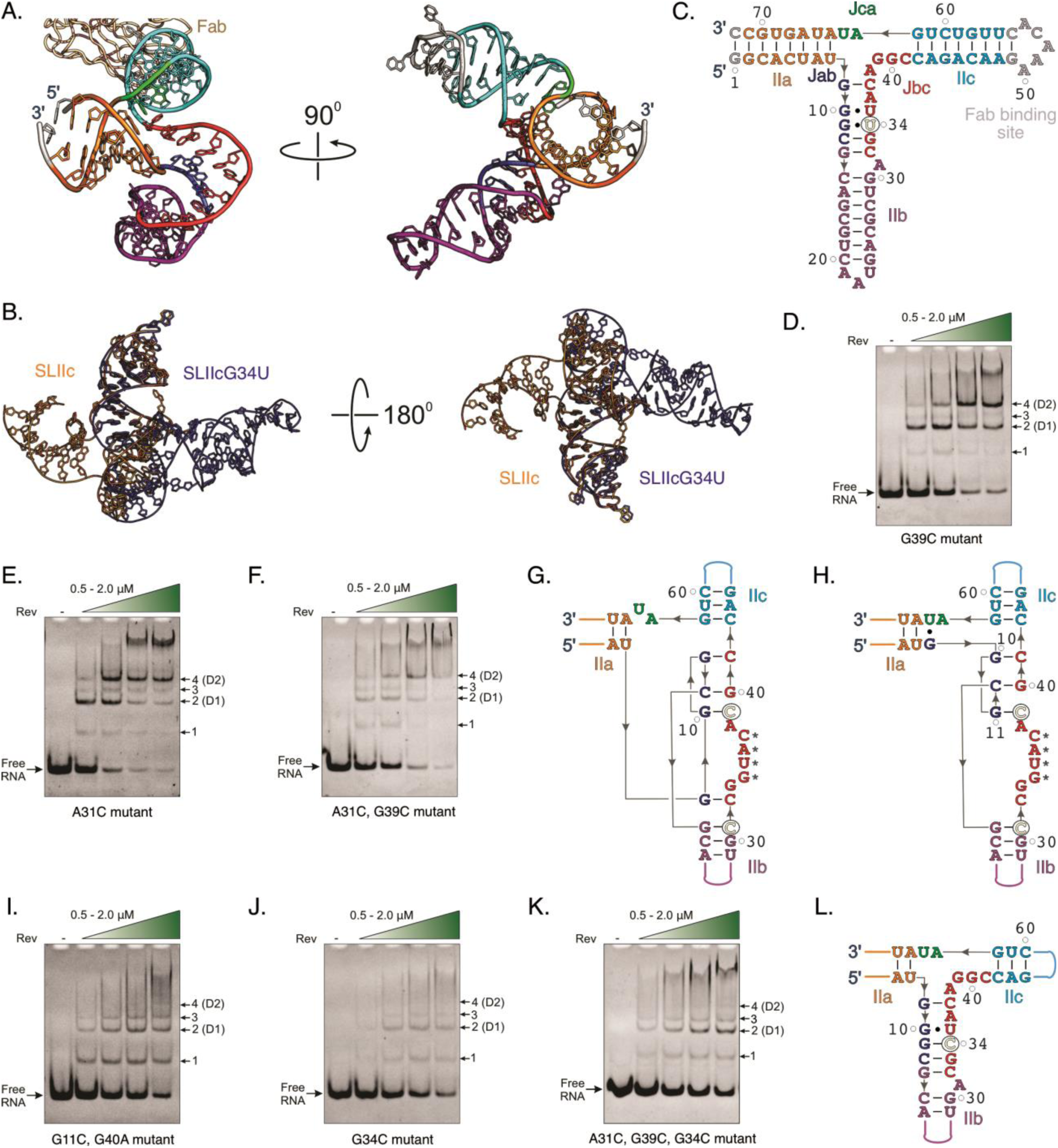
Crystal structures of the SLII mutants in alternative conformations. (A) The crystal structure of the G34U mutant construct cocrystallized with the Fab and determined at 3.0 Å resolution. (B) Aligning the IIc stem-loops of the SLIIc (orange) and SLIIcG34U (blue) shows relatively more compact SLIIc and extended G34U crystal structures. (C) The crystal-derived secondary structure for the G34U. For clarity, the Fab is obscured in the rotated views in panels A and B. All figure panels and corresponding labels, if any, are colored analogously for facile comparison. nPAGE showing the Rev-binding and oligomerization with the compact-fold-stabilizing mutants (D) G39C, (E) A31C, and (F) A31C, G39C. (G) The crystal-derived secondary structure for A31C (3.1 Å resolution) and double mutant A31C, G39C (2.75 Å resolution). (H) The structure of one of the symmetry-mates for the A31C, G39C mutant with a slightly different configuration of the 3WJ. nPAGE showing the Rev-binding and oligomerization with the extended-fold-stabilizing mutants (I) G11C, G40A (J) G34C, and (K) A31C, G39C, G34C. (L) The crystal-derived secondary structure for G34C (3.0 Å resolution). For the gels, the green color-gradient triangle shows the Rev concentration ranging from 0.5 to 2.0 µM with the 0.5 µM increment in each lane. The mutated nucleotides are colored yellow and circled for clarity. All figure panels and corresponding labels, if any, are colored analogously for facile comparison.

### Structure-guided mutations favor alternative SLII conformations

Based on the observed tendency of the SLII to adopt multiple conformations, it is plausible to assume that these conformations coexist in solution, and Rev-induced structural rearrangements may modulate the fate of Rev binding and oligomerization, thus affecting the nuclear export of HIV-1 RNAs. To test this hypothesis, we introduced mutations designed to favor either the compact conformation (like the non-mutant structure) or the extended conformation (like the G34U mutant structure) of the SLII and conducted the Rev binding assays. First, we incorporated mutations designed to stabilize the compact conformation by introducing A31C and G39C mutations that convert the non-canonical purine: purine pairs G12•A31 and G10•G39 into the canonical pairs G12:C31 and G10:C39 in the compact conformation (Figure 4D-F), without impacting the base-pairing pattern in the G34U-like extended conformation (Figures 1D, 4C). Interestingly, these mutations generally showed similar Rev binding patterns as the non-mutant SLIIc (compact form) and perhaps facilitated the Rev oligomerization cooperativity, suggesting that this conformational state of the SLII allows cooperative Rev binding and oligomerization. These results also imply that G34-C37 nucleotides observed in crystal contacts constitute the Rev-binding site, and the compact form perhaps represents a Rev-bound state captured in the crystal due to stabilization by the crystal contacts, which is consistent with what has been referred to in previous studies as an “excited state” of the SLII that is favorable for Rev oligomerization ^38^. Consistent with gel results, crystal structures of the A31C and A31C, G39C mutants (Figures 4G, S15, S16 and see Table S1 for statistics) showed identical folds as the non-mutant, indicating that the conformational state of the SLII has more pronounced effect on Rev binding and multimerization than the specific nucleotide identity. Notably, one of the SLII molecules within the crystallographic asymmetric unit for the A31C, G39C double mutant had a slightly different rearrangement of the base-pairing pattern compared to the other three molecules (Figures 4H, S16), underscoring the high plasticity of this SLII 3WJ.

Next, we populated the extended conformation and introduced G11C, G40A, and G34C mutations that perturbed the base-pairing patterns in the compact fold without much effect on the extended fold. These mutants showed decreased Rev binding affinity, and dimerization was mainly limited to one of the sites, presumably the D1 site (Figures 4I, J). Such deleterious effects on Rev binding were not rescued by compact fold-stabilizing of A31C and G39C mutations in the context of G34C mutation (Figure 4K), indicating that the extended fold perhaps represents the most thermodynamically stable conformation of SLII, which is consistent with structural similarities between solution NMR ^37^ and crystal structures of the extended IIb^35–37^. As expected, the crystal structure of the G34C mutant exhibited an identical fold as the G34U mutant (Figures 4L, S17 and see Table S1 for statistics), supporting that the G34 nucleotide is vital to switching the conformation of the SLII. Moreover, mutation of G34 to A34 to maintain the non-canonical base-pairing displays a similar Rev-binding pattern as the non-mutant form (Figure 5A), indicating that the purine-purine mismatch-pair maintains the plasticity and dynamicity of the 3WJ structure to enable Rev binding. Additionally, the insertion of the U14 nucleotide, relatively distant from the 3WJ site, showed a similar pattern of Rev binding as the non-mutant (Figure 5B), further supporting that the Rev-binding and oligomerization occur within the 3WJ. Moreover, the binding pattern of non-mutant SLII seems to be somewhat in between compact and extended forms, indicating that SLII can exist in multiple conformations in equilibrium in the solution, and structural rearrangements upon Rev binding within SLII may define the fate of the Rev binding and oligomerization.

**Figure 5.**
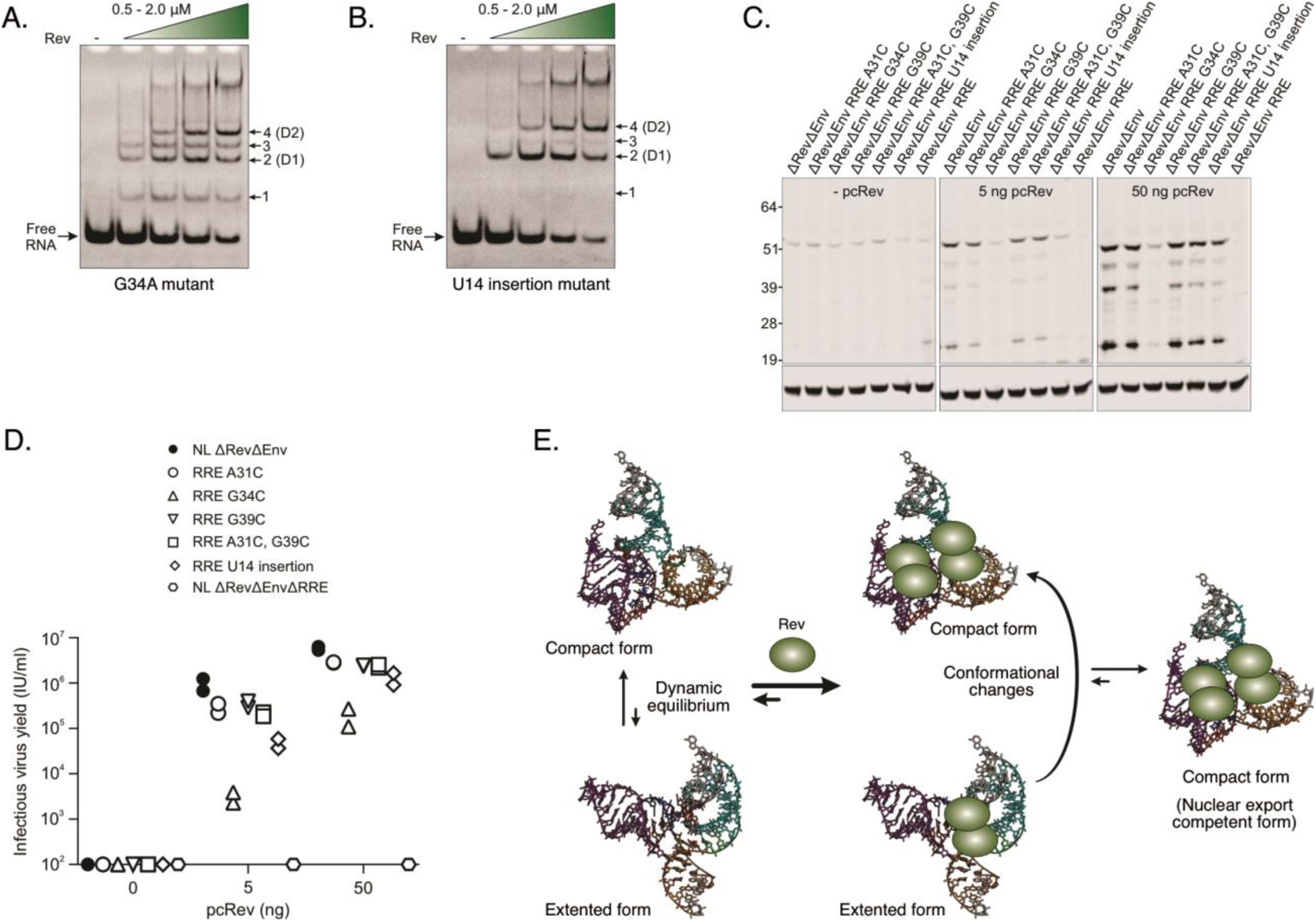
A structure-based model for RRE-mediated nuclear export of HIV-1 RNA. nPAGE showing the Rev-binding and oligomerization with (A) G34A mutant and (B) U14 insertion. The green color-gradient triangle shows the Rev concentration range from 0.5 to 2.0 µM with the 0.5 µM increment in each lane. (C) Gag expression and (D) infectious virion yield following transfection of HEK 293T cells with HIV-1 ΔRevΔEnv proviral plasmids, along with an Env-complementing VSV-G expression plasmid and 0 ng, 5ng or 50ng of a Rev expression plasmid. The HIV-1 Gag expression was assessed by western blotting using an HSP90 loading control. The infectious virion yield was determined by inoculating MT4 LTR-GFP indicator cells with transfected cell supernatants. (E) A proposed model for Rev-RRE-mediated nuclear export of HIV-1 RNAs. The model highlights the high structural plasticity of HIV-1 RRE SLII that enables it to adopt multiple conformations to modulate Rev binding and oligomerization.

### Conformational changes correlate well with Rev activity in cells

We performed cellular assays to test how alternative folds of the SLII construct observed in crystal structures modulate the Rev activity and nuclear export of HIV-1 RNAs. In these assays, we measured Gag protein expression and infectious virion yield following transfection of HEK 293T cells with an HIV-1 ΔRevΔEnv proviral plasmid, an Env-complementing VSV-G and a Rev expression plasmid. The Gag expression was assessed by western blotting using HSP90 protein as a loading control. The infectious virion yield was determined by inoculating MT4 LTR-GFP indicator cells with transfected cell supernatants. Whereas A31C, G39C, and double mutant A31C, G39C behaved similarly to the wild-type construct, the Gag expression and infectious virion yield were significantly reduced for the G34C mutant (Figures 5C, D). The U14 insertion mutant moderately reduced Gag expression and infectious virion yield (Figures 5C, D). The lower Rev activity and nuclear export of HIV-1 RNAs, as indicated by the reduced Gag expression and infectious virion yield, is consistent with stabilizing the extended fold of the SLII as observed in our crystal structures. Notably, we observed no significant reduction in Gag expression and infectious virion yield for the compact fold-stabilizing mutants (Figure 5C, D), suggesting that the compact crystal structure observed for the non-mutant likely represents a Rev-bound form, and the crystal contacts in crystal lattice perhaps coincidently stabilized such a fold. Consistent with the previous crystal structure of the IIb with a Rev dimer ^37^, several biochemical studies ^22, 23, 31^ indicated that the flexibility at the 3WJ orients the Rev molecules for adaptive recognition, binding affinity and cooperativity. NMR studies with the isolated IIb constructs^39^ also showed such flexibility in the purine-rich region of SLII. Our proposed mechanism of HIV-1 nuclear export not only explains previous observations pertinent to the RRE structure-mediated Rev binding and oligomerization, but it also supports a growing view that RNAs form multiple structures that can exist as a dynamic equilibrium of alternative conformations with several secondary structures, modulating the biological activity ^38^. For HIV-1 RRE, propose that SLII may exist in a dynamic equilibrium of conformations toggling between a compact and extended form, and binding the Rev shifts the equilibrium toward the compact conformation, facilitating cooperative Rev binding and oligomerization, thus allowing robust interactions with cellular nuclear export machinery.

## DISCUSSION

Based on the observation of multiple crystal structures of the full-length SLII in two distinct conformations and Rev binding studies with the wild-type and mutant SLII constructs, we propose that the structural plasticity and conformational dynamics of the HIV-1 RRE SLII regulate Rev binding and oligomerization to define the fate of HIV-1 RNA nuclear export. As illustrated in Figure 5E, in this model, the SLII can exist in at least two forms in solution – the compact and extended conformations. The extended conformation is consistent with previous NMR studies, and it may represent a thermodynamically more stable fold in solution in the absence of Rev. The binding of Rev to SLII thus drives the overall conformational equilibrium towards the compact form, with each subsequent Rev binding event reinforcing this shift in equilibrium towards a compact structure. Our studies suggest a model where the compact form of the SLII allows cooperative binding and multimerization of Rev within the HIV-1 RRE, leading to downstream interactions of the Rev-RRE complex with the host’s nuclear export machinery. It is critical to note that wild-type SLII structure in solution may assume different 3WJ configurations, and mutational effects observed in our Rev-binding and cellular activity measurements may be a combined effect of the disruption of specific interactions between SLII and Rev rather than the solo effect arising from toggling of RNA conformational equilibrium. However, observing multiple crystal structures for this RNA within the same crystallographic asymmetric unit and in independent crystals strongly argues that the structural plasticity of HIV-1 RRE is a significant factor in modulating Rev oligomerization. Our ongoing studies of full-length RRE and RRE-Rev complex structures aim to validate these structural insights into HIV-1 RNA nuclear export mechanisms.

Notably, the mechanism of Rev-SLII interactions based on our SLII structure appears to be quite different than what was proposed based on the crystal structure of a Rev dimer in a complex with a synthetic IIb hairpin construct (PDB: 4PMI) ^37^. This model predicted a series of non-canonical G•A and G•G pairs as a part of the 3WJ and stem IIb that comprised two high-affinity Rev binding and dimerization sites. However, consistent with biochemical probing and SHAPE studies ^20, 24, 29–31^, the nucleotides interacting with Rev are an integral part of 3WJ in our structures rather than an IIb bulge, suggesting that multiple high-affinity Rev binding sites are incorporated within the SLII 3WJ. Another proposed model for RRE-Rev binding was derived from the results of a SAXS-based study by Fang *et al*. ^39^ using an intact 232-nt HIV-1 RRE construct. Starting from a SAXS-derived A-shaped structure of the RRE, this work suggested two Rev binding sites located within the SLII and SLI and separated by about 55 Å. As the spacing between the two RNA-binding domains in the Rev dimers resembles the spacing between these two binding sites, the authors hypothesized that initial Rev binding helps to bridge these two Rev binding sites. Although the high-resolution tertiary structure of the intact RRE has yet to be determined, our results suggest that, unlike the SAXS-based model, SLII alone can form an oligomeric RRE-Rev complex, which is also consistent with a recent native mass-spectrometry-based study that showed SLII can bind up to five Rev molecules ^50^. Since the intact RRE has been shown to bind up to eight Rev molecules ^18, 51, 52^ and SLII contains multiple Rev dimerization sites, SLI may serve as a secondary Rev dimerization site that facilitates the connection between SLII and SLI sites through Rev-Rev interactions. Our initial results indicate a single Rev dimer binding to an SLI construct (Figure S18). However, it is important to note that SL1 may create an initial dimer binding site or promote the proper arrangement of the RRE-Rev oligomeric complex within the intact RRE, which warrants further investigation.

While preparing this manuscript, Tipo *et al*. ^40^ published a 2.85 Å resolution crystal structure (PDB: 8UO6) of the SLII fused with a tRNA scaffold within the stem IIa ^40^. While some conclusions drawn from their study, such as the location of the high-affinity Rev-binding sites within the 3WJ rather than the stem IIb and the presence of the higher and lower-affinity binding sites for Rev dimerization are consistent with our study, their SLII structure itself, especially around the 3WJ, is quite different than ours. Their 3WJ junction appeared extended and unstructured, with most junction nucleotides unpaired. However, our SLII crystal structures have a well-organized 3WJ with extensive canonical and non-canonical base-pairing interactions between the junction nucleotides. They proposed closed and open conformations of the SLII that differ in reordering a couple of non-canonical pairs observed in the two molecules within the asymmetric unit of the same crystal. However, the impact of extensive crystal contacts around the 3WJ for stabilizing the open or closed conformation was unclear. Notably, the unpaired nucleotides involved in crystal contacts in one of our structures (G34 to C37) also remained unpaired in their structure, matching closely with our molecular dynamics results. Our G34U and G34C mutants also adopted a distinct fold compared to those conformations, although this G34U mutation seems to substitute a non-canonical G10•G34 pair by G10•U34 that would have further stabilized their structure. These substantial differences in the 3WJ architecture extend to the overall positioning of the helices in these structures. Unlike the IIa and IIb stems coaxially aligned and the IIc is almost perpendicular to the IIa/IIb junction axis in their structure, our extended conformation shows IIa and IIc stems coaxially aligned and IIb protruding out almost perpendicularly from the IIa/IIc junction axis. Overall, these structures highlight the tendency of HIV-1 RRE SLII to adopt multiple conformations, supporting a model where the structural plasticity of the RNA is an important mechanism regulating RRE-Rev interactions and function.

The nuclear export of the intron-containing RNAs is essential for HIV-1 replication, so the RRE-Rev platform has been an attractive therapeutic target ^53, 54^. While the structures of isolated IIb have been used to direct the development of small-molecule drugs against HIV-1 infections ^53, 54^, our structural studies with intact SLII provide better opportunities to screen for or structure-based design of the inhibitors that target the well-structured high-affinity Rev binding sites within the SLII 3WJ. Moreover, the structural plasticity of SLII observed in multiple high-resolution crystal structures would make it possible to develop small molecule drugs that selectively bind one of the conformations, especially those that significantly reduce nuclear export of HIV-1 RRE, offering various paths toward the development of anti-HIV therapeutics relative to the approaches that target a single specific conformation of the RRE.

## METHODS

### Materials

The sequences of the RRE SLII constructs used in this study are listed in Table S2. Unless stated otherwise, all ssDNA templates and PCR primers for these constructs were purchased from Integrated DNA Technologies (IDT) Inc.

### RNA synthesis and purification

The RNA constructs for this study were synthesized by *in vitro* transcription. DNA template with a T7 promoter sequence for transcription reaction was produced by PCR amplification of ssDNA purchased from IDT. The first two nucleotides of reverse primer were 2′ OMe modified to reduce the 3′ end heterogeneity of the transcript ^55^. The transcription reaction was conducted for 3 hours at 37 °C in a buffer containing 40 mM Tris-HCl, pH 8.0, 2 mM spermidine, 10 mM NaCl, 25 mM MgCl_2_, 0.1 mM EDTA, 1 mM DTT, 40 U/ml RNase inhibitor, 5 U/ml TIPPase, 5 mM of each NTP, 50 pmol/ml DNA template, and 50 μg/ml homemade T7 RNA polymerase ^56^. The reaction was then quenched by adding 10 U/ml DNase I (Promega) and incubating at 37°C for 1 hour. All RNA samples were purified by denaturing polyacrylamide gel electrophoresis (dPAGE). The RNA band was visualized by UV shadowing, excised from the gel, crushed, and eluted overnight at 4 °C in 10 mM Tris, pH 8.0, 2 mM EDTA, and 300 mM NaCl. The buffer of eluted RNA was exchanged with pure water three times using a 10 kDa cut-off Amicon column (Millipore Sigma). RNA was collected, aliquoted into 300 μl fractions, and stored at −80°C until further use.

### Rev expression and purification

The HIV-1 Rev gene was cloned into a pET19b vector (GenScript) with a 6x-His tag at the N-terminal and MBP tag, and the plasmid containing this MBP tag-fused Rev was transformed into BL21 (DE3) E. coli. The bacterial culture was grown into LB supplemented with 100 µg/ml of ampicillin at 37 ℃ with 250 rpm in a 5 ml starter culture for ∼ 12 hours (overnight), which was then used to inoculate 1L culture for overnight growth (until the OD of ∼ 0.6). The protein overexpression was induced using IPTG to the final concentration of 0.5 mM for ∼12 hours (overnight) at 18 ℃ before harvesting the bacteria through centrifugation at 6000 g for 20 minutes at 4 ℃. The cell pellets were then resuspended using lysis buffer (50 mM Tris-base, pH 7.5, 200 mM NaCl and 5% glycerol) and lysed using sonication. The lysate was centrifuged at 13000 rpm at 4°C, and the supernatant was passed through a 0.45-micron filter. The clarified lysate was then applied to a 5 ml MBPTrap™ HP column (Cytiva). The protein was eluted from the column isocratically with a buffer (25 mM Tris-base, pH 7.5, 200 mM NaCl, 5% glycerol, 1 mM TCEP, and 10 mM maltose monohydrate) after washing the column with 5 column volumes of the lysis buffer. The protein was subsequently applied to a 5 mL HiTrap™ Heparin HP column (Cytiva) and eluted using a percentage gradient of a buffer containing 40 mM Tris-base, pH 7.5, 2 M NaCl, and 5 mM TCEP. The eluted fractions were collected and further purified using size-exclusion chromatography with a HiLoad® 26/60 Superdex® 75 pg column (Cytiva) and a buffer composed of 20 mM Tris-base, pH 7.5, 140 mM KCl, 5 mM TCEP, and 1 mM MgCl_2_. The single-peak protein fractions were pooled, concentrated using Amicon centrifugal filters (molecular weight cut-off 10 kDa, Millipore Sigma) and stored at −80°C.

### Fab expression and purification

The Fab BL3-6 expression plasmid was a kind gift from Joseph Piccirilli, the University of Chicago. The Fab was expressed and purified according to published protocols ^57–59^. Briefly, the plasmid was transformed into 55244 *E. coli* competent cells and streaked into an LB-agar plate with 100 µg/ml of carbenicillin. Several colonies were selected to inoculate a 15 ml starter culture and grown at 30°C for 8 hours. The starter culture was then used to inoculate 1 liter of 2xYT media, and cells were grown for 24 hours at 30°C. For Fab overexpression, the cells were centrifuged at 22°C and 6000g for 10 minutes, resuspended in 1-liter phosphate-depleted media, and grown for 24 hours at 30°C. The cells were harvested by centrifugation at 4°C and 6000g for 10 minutes, resuspended in PBS, pH 7.4 buffer with 0.01 mg/ml bovine pancreas DNase I (Sigma-Aldrich), and 400 mM Phenylmethylsulfonyl fluoride (PMSF), and lysed by sonication (Qsonica, Cole-Parmer). The mixture was first centrifuged at 18000 rpm, the clear lysate was filtered through a 0.45-micron filter (VWR), and the Fab was purified using the Bio-Rad NGC fast protein liquid chromatography (FPLC) system. First, the lysate was passed through a Hi-trap protein A column (Cytiva), and the captured Fab was eluted with 0.1 M acetic acid. The fractions were collected, diluted using PBS pH 7.4 buffer, and loaded into a Hi-trap protein G column (Cytiva). The eluted Fab fractions from the protein G column in 0.1 M glycine, pH 2.7, were collected, diluted with a 50 mM NaOAc, 50 mM NaCl, pH 5.5 buffer, and loaded into a Hi-trap heparin column (Cytiva). Finally, the Fab fractions eluted from the heparin column by the gradient elution using 50 mM NaOAc, 2 M NaCl, and pH 5.5 buffer were collected, and buffer was exchanged 3 times with PBS pH 7.4 using 30 kDa cut-off Amicon column (Millipore Sigma). The concentrated Fab was collected, analyzed by 12% SDS-PAGE, and tested for RNase activity using the RNaseAlert kit (Ambion, www.thermofisher.com). The aliquots (∼300 μl) of purified Fab were stored at −80 °C.

### Native gel electrophoresis assay

For Fab-binding with crystallization constructs, about 200 ng RNA (∼ 0.5 μM) in water was refolded in a buffer containing 10 mM Tris-HCl, pH 7.4, 5 mM MgCl2, and 300 mM NaCl. For this, RNA was heated at 90°C for 1 minute, and an appropriate volume of the refolding buffer was added, followed by incubation at 50°C for 10 minutes and in ice for 5 minutes. The refolded RNA was then incubated for 30 minutes at room temperature with different Fab. The protein-RNA complex samples were mixed with an appropriate volume of native gel loading solution that contained 30% glycerol, 0.1% bromophenol blue, and xylene cyanol. These samples were loaded onto 10% native polyacrylamide gel and run at 115 V in 0.5× TBE buffer (50 mM Tris-base, 50 mM boric acid, and 1 mM EDTA, pH 7.5). For Rev-binding with SLII constructs, 150 ng of RNA was incubated with 1× refolding buffer containing 10 mM Tris-HCl, pH 7.5, 5 mM MgCl2, and 10 mM NaCl in the presence of Rev protein at 37 °C for 1 hr. The samples were loaded onto 5% native polyacrylamide gel and run at 100 V in 1× TB buffer (50 mM Tris-base and 50 mM boric acid, pH 7.5). The native gels were stained with ethidium bromide and imaged using the Azure 200 gel documentation system (Azure Biosystems).

### Crystallization

The RNA sample (∼ 600 μg) was refolded in a refolding buffer containing 10 mM Tris-HCl, pH 7.5, 5 mM MgCl_2_, and 300 mM NaCl as described above for the native gel electrophoresis assay. The refolded RNA was then incubated for 30 min at RT with 1.1 equivalents of the Fab and concentrated to 6 mg/ml using a 10-kDa cut-off, Amicon Ultra-1 column (Millipore Sigma). Then, Fab−RNA complexes were passed through 0.2 μm cut-off Millipore centrifugal filter units (www.emdmillipore.com). The Xtal3 Mosquito liquid handling robot (TTP Labtech, ttplabtech.com) was used to set up sitting-drop vapor-diffusion crystallization screens at room temperature (22°C) within a humidity-controlled (70%) chamber using commercially available screening kits from Hampton Research. Out of 672 conditions screened for each complex, the crystal hits were observed only for the SLIIc, SLIIcG34U and SLIIcA36U mutants in complex with the Fab BL3-6. However, SLIIcA36U-Fab complex crystals did not diffract to a high enough resolution to advance further to the structure-solving process. The best diffracting crystals for the SLIIc and SLIIcG34U complexes were observed within a week in several conditions. Drops containing suitable crystals were brought up to 40% glycerol for cryoprotection without changing the other compositions. The crystals were immediately flash-frozen in liquid nitrogen after being fished in the loops from the drops and shipped to the Brookhaven National Laboratory (BNL) for X-ray diffraction screening and data collection.

### Crystallographic data collection, processing, and analysis

The X-ray diffraction data sets were collected at the Brookhaven National Laboratory, NSLSII beamlines 17-ID-1 (AMX) and 17-ID-2 (FMX). Datasets were collected for several single crystals; however, the crystal grown in a condition with 0.2 M sodium acetate trihydrate, pH 7.0 with 20% PEG 3350 provided the best resolution of 2.42 Å for the SLIIc-Fab complex. The best diffracting crystal (3.0 Å resolution) for the SLIIcG34U-Fab complex was obtained in a condition with 0.2 M NaCl, 0.1 M Tris pH 8.0, 12.5% Polyvinylpyrrolidone, 10% PEG 4000. Likewise, the best diffracting crystal for the SLIIcG34C-Fab complex (3.0 Å resolution), SLIICA31C-Fab complex (3.10 Å resolution) and SLIIcA31CG39C-Fab complex (2.75 Å resolution) were obtained in 0.02 M citric acid, 0.08 M BIS-TRIS propane pH 8.8, 16% w/v polyethylene glycol 3350 and 0.03% ethanol; 0.2 M ammonium nitrate, 20% w/v polyethylene glycol 3350 and pH 6.2; and 0.2 M ammonium citrate dibasic, 20% w/v polyethylene glycol 3350 and pH 5.1, respectively. All the datasets were then integrated and scaled using its on-site automated programs (AutoProc). Some datasets were also processed and analyzed using the Xia2/Dials platform on the CCP4 ^60, 61^. The crystallographic data suggested that the SLIIc-Fab complex was crystallized in the P1 space group and contained two complexes within the crystallographic asymmetric unit (a = 72.36 Å, b = 76.24 Å, c = 82.60 Å, α = 116.56°, β = 94.91°, and γ = 102.75°) with the 55.72% solvent content (see Supplementary Table S1 for details of the data collection statistics). Unlike the SLIIc-Fab complex, the SLIIcG34U-Fab complex crystallized in the I222 space group, and it contained a single complex within the crystallographic asymmetric unit (a = 90.88 Å, b = 105.01 Å, c = 186.60 Å, α = 90°, β = 90°, and γ = 90°). Similarly, the SLIIcA31C-Fab complex was crystallized in the P1 space group. It contained two complexes within the crystallographic asymmetric unit (a = 72.87 Å, b = 75.87 Å, c = 83.51 Å, α = 63.17°, β = 71.85°, and γ = 77.75°). The SLIIcA31CG39C-Fab complex was crystallized in the P1 space group and contained four complexes within the crystallographic asymmetric unit (a = 84.37 Å, b = 95.64 Å, c = 111.24 Å, α = 77.486°, β = 76.34°, and γ = 74.25°). The SLIIcG34C-Fab complex crystallized in the I222 space group, and it contained a single complex within the crystallographic asymmetric unit (a = 90.78 Å, b = 106.63 Å, c = 187.33 Å, α = 90°, β = 90°, and γ = 90°). Details of data collection statistics are provided in Table S1.

For solving both structures, the initial phases were obtained by molecular replacement with the previously reported structure of Fab BL3-6 (PDB code: 8T29) ^43^ as the search model using Phaser on Phenix ^62^. Iterative model building and refinement were performed using the COOT ^63^ and the Phenix package ^62^. The RNA structure was built unambiguously by modeling the individual nucleotides into the electron density map obtained from the molecular replacement. The refinement used default NCS options and auto-selected TLS parameters in Phenix. The structure-related figures were made in PyMOL (The PyMOL Molecular Graphics System, Version 2.0 Schrödinger, LLC), and the figure labels were edited in CorelDraw (Corel Corporation, http://www.corel.com).

### Electron microscopy data collection, processing, and analysis

Samples for negative stain were prepared as previously described (Ohi: PMC389902) ^64^. The SLIIc-Fab complex sample from the crystallographic experiments was serially diluted in the corresponding buffer (10 mM Tris-HCl, pH 7.5, 5 mM MgCl_2_, and 300 mM NaCl). Formvar (10 nm)/carbon (1 nm)-coated copper grids with 400 mesh size (EMS FCF400-Cu-50) were glow discharged for 30 seconds with a 5mA current. 3.5µL of the sample was applied to the grid and incubated for 1 minute. The grid was then washed two times using two drops of water and then two drops of 0.75% uranyl formate. The grids were screened for ideal concentration on a Thermo Fisher Morgagni 100 kV TEM with Gatan Orius SC200 CCD Detector at 22,000 times magnification. A 1.2 mg/ml stock of the SLIIc-Fab complex was diluted 1:1000 and 1:5000 in sample buffer and imaged for data processing on a Thermo Fisher Tecnai T12 with Gatan Rio9 CMOS Detector at 36,000 times magnification corresponding to a size of 2.08Å per pixel was used for square targeting and data collection. Micrographs were collected on T12 using SerialEM^65^, imported into, and processed in cryoSPARC (v4.5.1) (PMID: 28165473) ^66^. CTF estimation was performed with CTFFIND4 ^67^. Particle picking was performed with crYOLO v1.9.6 using a general negative stain model ^68^. The box size was 200 pixels (i.e., 416 Å). About 800,357 particles for the 1:1000 dilution data set were subjected to two rounds of 2D classification, designating 100 and 50 classes. About 96912 particles for the 1:5000 dilution data set were subjected to one round of 2D classification designating the 50 classes.

### Molecular dynamics simulation

The molecular dynamics simulations were conducted departing from the crystal structure of the SLIIc construct. For the A63G system, guanine was modeled onto the adenine at position 63 of the SLIIc crystal structure. All MD simulations were carried out using AMBER22 ^46^. The systems were neutralized with Na^+,^ and additional ions were added to reach a bulk concentration of 0.14 M NaCl in a truncated octahedron box with 88 Å real-space lattice vector lengths. The systems were built with ff99OL3 RNA ^69^ force field, TIP4P/Ew water model ^70^ and corresponding Na^+^ and Cl^-^ ion parameters ^71^. The simulations were then performed under periodic boundary conditions at 298 K using a 12Å nonbonded cutoff and PME electrostatics ^72, 73^. The Langevin thermostat with γ = 5 ps^-1^ and Berendsen isotropic barostat with τ = 1 ps were used to maintain constant pressure and temperature. A 1 fs time step was used along with the SHAKE algorithm to fix hydrogen bond lengths ^74^, with three-point SHAKE for the water molecules ^75^. As reported, the simulation boxes were equilibrated stepwise ^76^, yielding ∼50 ns of simulation time. All production simulations were carried out in constant pressure and temperature (NPT) ensembles, and 1250 ns of aggregate production were achieved per system. Root mean square deviations (RMSD) were calculated to quantify the structural changes and identify robust versus flexible molecule regions throughout the simulation, using the starting structures as the reference.

### Cellular Rev-activity measurements

The HIV-1NL4-3 ΔEnv, HIV-1NL4-3 ΔRevΔEnv, HIV-1NL4-3 ΔRevΔEnvΔRRE, pcRev, and pVSV-G have been described previously ^77–79^. Point mutations (A31C, G34C, G39C, A31C/G39C) and the insertion of U at position 14 in RRE SLII were made to the corresponding residues of HIV-1NL4-3 ΔRevΔEnv through overlap extension PCR, using 5′ and 3′ oligonucleotides flanking unique NheI and XhoI restriction sites in combination with oligonucleotides containing the point mutations or insertion. PCR products were digested and ligated into HIV-1NL4-3 ΔRevΔEnv using NheI and XhoI. The following oligonucleotides (Integrated DNA Technologies, Inc) were used to obtain the RRE SLII mutant HIV-1NL4-3 ΔRevΔEnv plasmid constructs. NheI: 5′ primer: CTG TAG TTA GCT AGC AAA TTA AGA GAA CAA; XhoI: 3′ primer: CTA GGT CTC GAG ATA CTG CTC CCA CCC CATC; A31C: 5′ primer: TAT GGG. CGC AGC GTC AAT GAC GCT GCC GGT ACA GGC CAG ACA ATT ATT GTC TGA TAT AGT; A31C: 5′ primer: ACT ATA TCA GAC AAT AAT TGT CTG GCC TGT ACC GGC AGC GTC ATT GAC GCT GCG CCC ATA; G34C: 5′ primer: TAT GGG CGC AGC GTC AAT GAC GCT GAC GCT ACA GGC CAG ACA ATT ATT GTC TGA TAT AGT; G34C: 3′ primer: ACT ATA TCA GAC AAT AAT TGT CTG GCC TGT AGC GTC AGC GTC ATT GAC GCT GCG CCC ATA; G39C: 5′ primer: TAT GGG CGC AGC GTC AAT GAC GCT GAC GGT ACA CGC CAG ACA ATT ATT GTC TGA TAT AGT; G39C: 3′ primer: ACT ATA TCA GAC AAT AAT TGT CTG GCG TGT ACC GTC AGC GTC ATT GAC GCT GCG CCC ATA; A31C, G39C: 5′ primer: CTT GGG AGC AGC AGG AAG CAC TAT GGG CGT CAG CGT CAA TGA CGC TGA CGG TAC AGG CCA; A31C, G39C: 3′ primer: TGG CCT GTA CCG TCA GCG TCA TTG ACG CTG ACG CCC ATA GTG CTT CCT GCT GCT CCC AAG. HEK 293T cells (ATCC Cat#CRL-3216) were maintained in Dulbecco’s Modified Eagle Medium DMEM (ThermoFisher Cat#11995065) with 10% fetal calf serum (FCS) (Sigma, Cat# F8067-500ML) and gentamicin (ThermoFisher Cat#15710064). MT4-LTR-GFP reporter cells ^80^ were maintained in Roswell Park Memorial Institute RPMI (ThermoFisher Cat#11875119) with 10% FCS, gentamicin, and 1.25 μg/ml puromycin (Sigma-Aldrich Cat# P8833-100MG).

About 150,000 HEK 293T cells were co-transfected with 500 ng HIV-1NL4-3 ΔEnv, HIV-1NL4-3 ΔRevΔEnv, HIV-1NL4-3 ΔRevΔEnvΔRRE, or the point mutants of RRE as indicated, 50 ng of pVSV-G, and varying amounts of pcRev (0, 5 or 50 ng) using 1 mg/ml polyethyleneimine with a DNA: PEI ratio of 1:4 (Polysciences Cat#23966). Forty-eight hours post-transfection virus supernatants were harvested and 0.22 µm filtered (Fisher Cat#09-720-3), and cells were lysed in 1× SDS sample buffer. To determine the viral titer, supernatants were 5-fold serially diluted in RPMI and added to 50,000 MT4-LTR-GFP reporter cells. Forty-eight 48 hours post-infection, MT4-LTR-GFP cells were fixed with a final concentration of 2% paraformaldehyde (Sigma Cat#P6148). GFP+ cells were enumerated by flow cytometry (Attune NxT, Thermo) and used to calculate the infectious units of virus per ml of supernatant. For western blotting, cell lysates were separated by electrophoresis on NuPage 4-12% Bis-Tris gels (Invitrogen Cat# NP0323BOX) and transferred onto nitrocellulose membranes (GE Healthcare Cat# 10600003). Membranes were probed with primary antibodies against HIV-1 CA p24 (1:100) (NIH AIDS Reagent Catalog #530) and HSP90 (1:3,000) (Proteintech Cat# 13171-1-AP), followed by secondary antibodies (1:10,000) Goat anti-Mouse IRDye 680RD (LiCor 926-68070) and Goat anti-Rabbit IRDye 800CW (LiCor 926-32211). Membranes were scanned on a LiCor Odyssey M (LiCor). Image files were analyzed and exported using Image Studio 5.2 (LiCor).

### Statistics and reproducibility

The native PAGE experiments for all RNA constructs were performed in triplicate, and each replicate produced similar results.

## DATA AVAILABILITY

The atomic coordinates and structure factors for the reported crystal structures of SLIIc, SLIIcG34U, SLIIcG34C, SLIIcA31C, and SLIIcA31C G39C, all in complex with Fab BL3-6 have been deposited with the Protein Data Bank under accession codes 9C2K, 9C75, 9E7G, 9E7D, and 9E7E, respectively. All data are available in the main text or supplementary materials. The authors will provide plasmids for the Fab BL3-6 and Rev expressions upon reasonable request. Requests should be directed to D.K.

## ACKNOWLEDGEMENTS

This work was mainly supported by the Center for Structural Biology of HIV-1 RNA (CRNA) Collaborative Development Pilot Grant Program (NIH NIAID1U54AI170660, Sub-award SUBK00019302) to D. K. and partly by the NIH T32 grant (GM066706) to M. O. and NIH GM062248 to D. M. Y. The crystallographic data was collected at the NSLSII beamlines (17-ID-1 and 17-ID-2) in Brookhaven National Laboratory (BNL) using the beamtime obtained through NECAT BAG proposal # 311950. The NIH-NIGMS primarily supports the BNL’s Center for Bio-Molecular Structure (CBMS) through a Center Core P30 Grant (P30GM133893) and by the DOE Office of Biological and Environmental Research (KP1607011). The NSLS2 is the U.S. D.O.E. Office of Science User Facility operated under Contract No. DE-SC0012704. Computational resources were provided by the Office of Advanced Research Computing (OARC) at Rutgers, The State University of New Jersey; the Advanced Cyberinfrastructure Coordination Ecosystem: Services and Support (ACCESS) program (supercomputer Expanse at SDSC through allocation CHE190067); and the Texas Advanced Computing Center (TACC) at the University of Texas at Austin (supercomputer Frontera through allocation CHE20002). The authors also thank Prof. Micheal F. Summers, University of Maryland, Baltimore County, for critical review and feedback on the initial manuscript.

## AUTHOR CONTRIBUTIONS

D.K. and M.O. conceived and designed the experiments. M.O. prepared the samples, conducted most of the crystallographic and binding experiments, and solved the crystal structures with the help of D.K. L. Hudson purified the Rev protein and helped M.O. with binding experiments under the mentorship of J.M. A. Photenhauer performed and analyzed the electron microscopy work in the laboratory of M.D.O. T. Zang conducted cellular Rev activity assays in the laboratory of P.D.B. L. Lerew & S.E. performed the molecular dynamics simulation study and analyzed the results with D.M.Y. Undergraduate researchers J.D., M.N. & H.P. assisted M.O. with biochemical experiments and crystallization trials set up. M.O. analyzed most of the biochemical and crystallographic data and interpreted the results with D.K. M. Ojha & D.K. wrote the manuscript with support from D.M.Y., M.D.O., J.M. & P.D.B. All authors reviewed the manuscript.

