## Supplementary Information for "Structural Plasticity of RRE Stem-Loop II Modulates Nuclear Export of HIV-1 RNA"

### Supplementary Figures

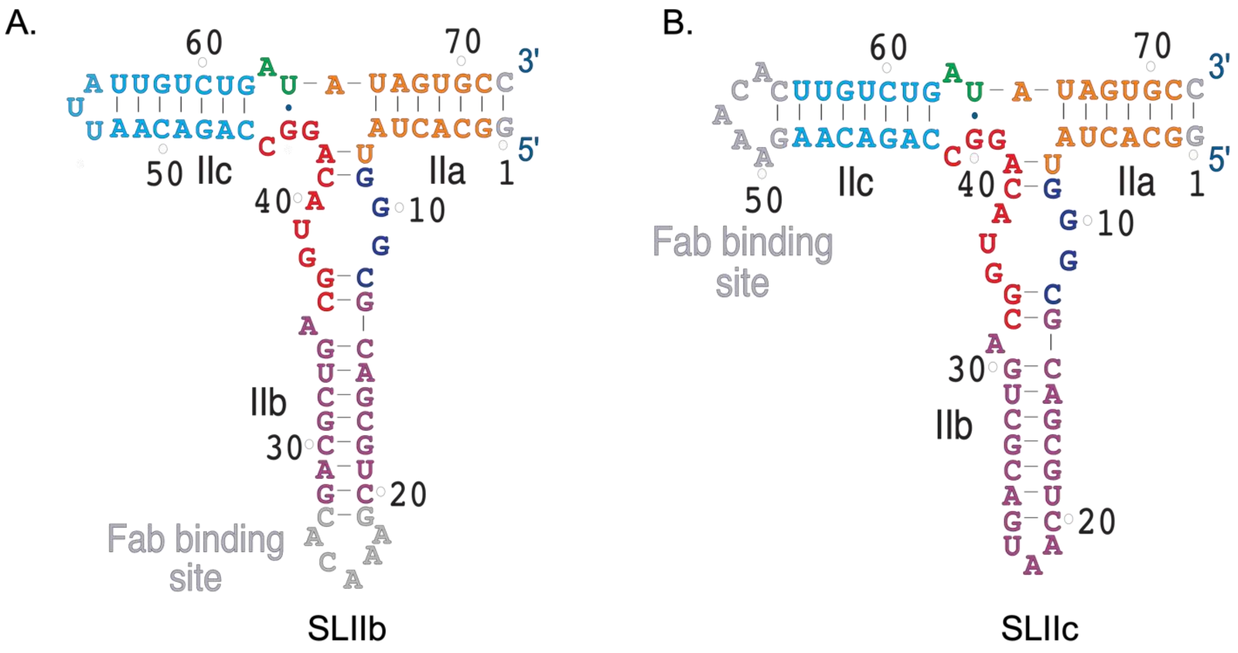


**Figure S1**. **Predicted secondary structures of the SLII crystallization constructs**. The 72-nt crystallization constructs with Fab-binding sequence grafted in the (A) IIb loop (SLIIb) and (B) IIc loop (SLIIc). The mutation or insertion in these crystallization constructs compared to the wild-type (Figure 1A), including the Fab binding sequence, are colored gray. The nucleotides are colored according to the crystal-derived secondary structure shown in Figure 1D.

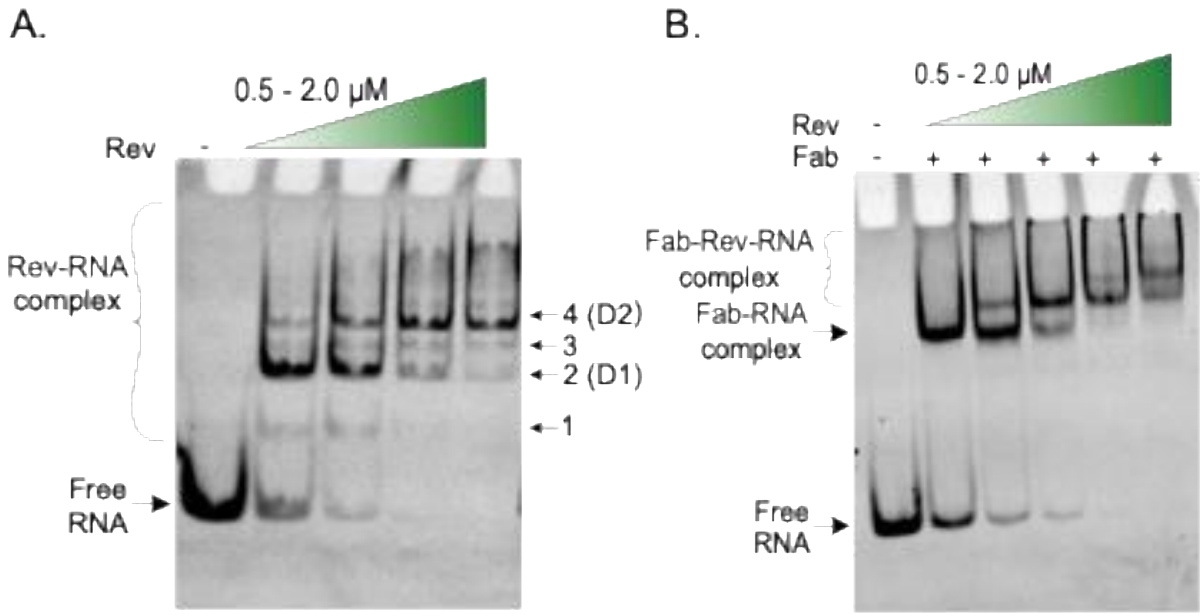


**Figure S2**. Native polyacrylamide gel electrophoresis (nPAGE) showing Rev binding with the SLIIb crystallization construct in the absence (A) and in the presence (B) of the Fab BL3-6. Each lane was loaded with ~ 150 ng (~ 0.5 µM) of RNA and 1.5 µM of the Fab. The Rev protein concentration was varied from 0.5 – 2 µM as indicated by the gradient-filled green triangles.

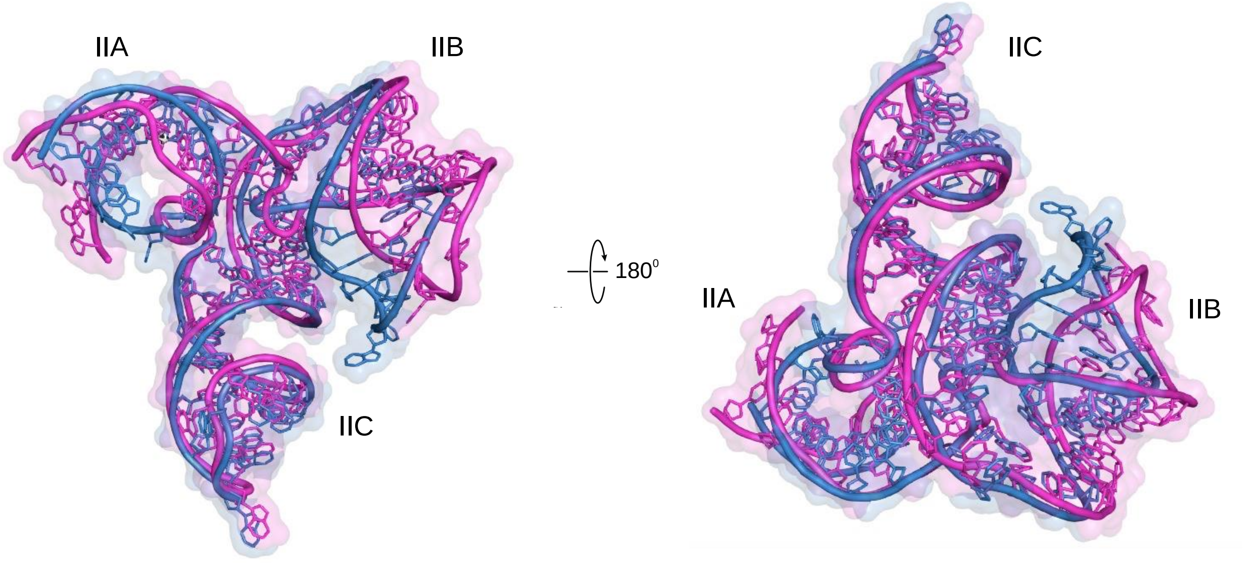


**Figure S3**. The superposition of the two SLIIc RNA molecules identified within the crystallographic asymmetric unit. The molecular surface is also shown for each molecule for facile comparisons. Overall, the two RNA molecules appear structurally similar (root mean square deviation, RMSD = 2.55 Å) with identical base-paring and stacking of nucleotides. However, we observed some deviations in relative orientations of the stems flanking the 3WJ, especially the stem-loop IIb, indicating the dynamic nature of the junction.

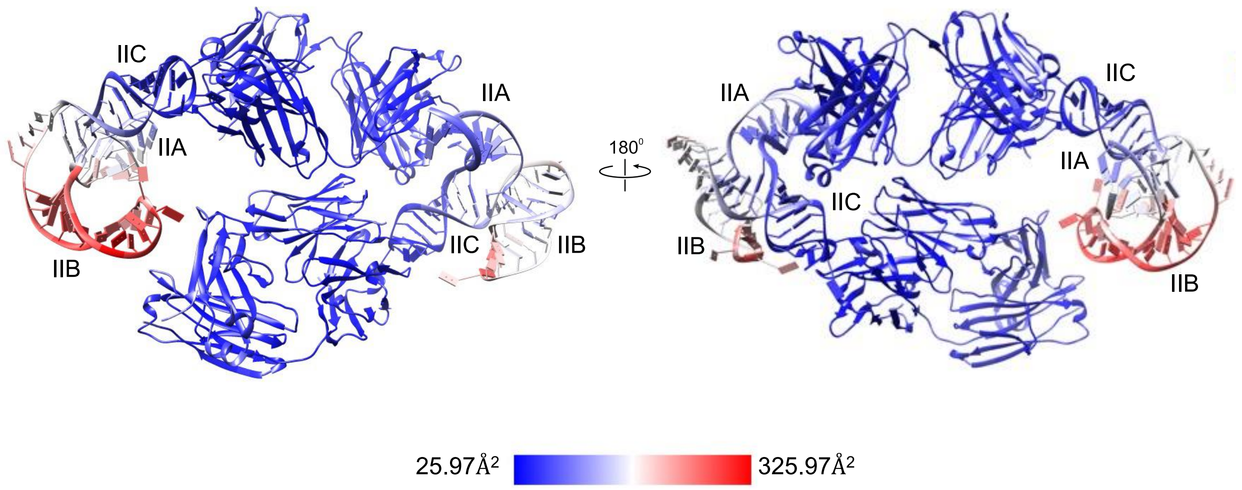


**Figure S4**. The crystal structure of the SLIIc in complex with Fab BL3-6 colored according to the crystallographic B-factors. Two molecules of the Fab-RNA complex observed in the crystallographic asymmetric unit are shown. The gradient from blue to red indicates the structure's lowest (25.97 $Å^2$) and the highest (325.97 $Å^2$) B-factors. Notably, the IIb stem-loop nucleotides showed the highest B-factors than those of IIa and IIc, indicating more flexibility of the IIb stem-loop compared to the IIa and IIc stem-loops.

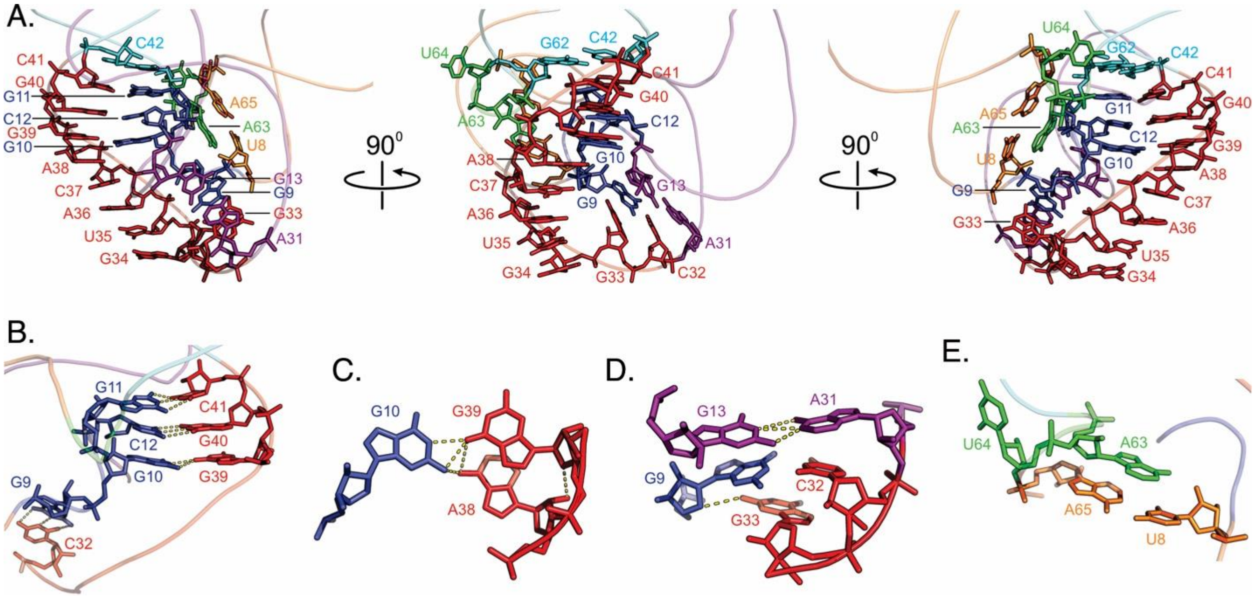


**Figure S5**. Interactions between nucleotides within the SLII 3WJ. (A) The overall structure of the SLIIc 3WJ. The specific interactions of Jab G9-C12 nucleotides with (B) the Jbc G39-C41 and (C) A38-G39 nucleotides. (D) The Jab G9 base pairs with C32 and interacts with the Jbc G33, where the G13:A31 non-canonical pair stacks from the top. (E) The specific interactions at or near the Jca. Yellow dashed lines in b-d indicate heteroatoms within hydrogen bonding distances.

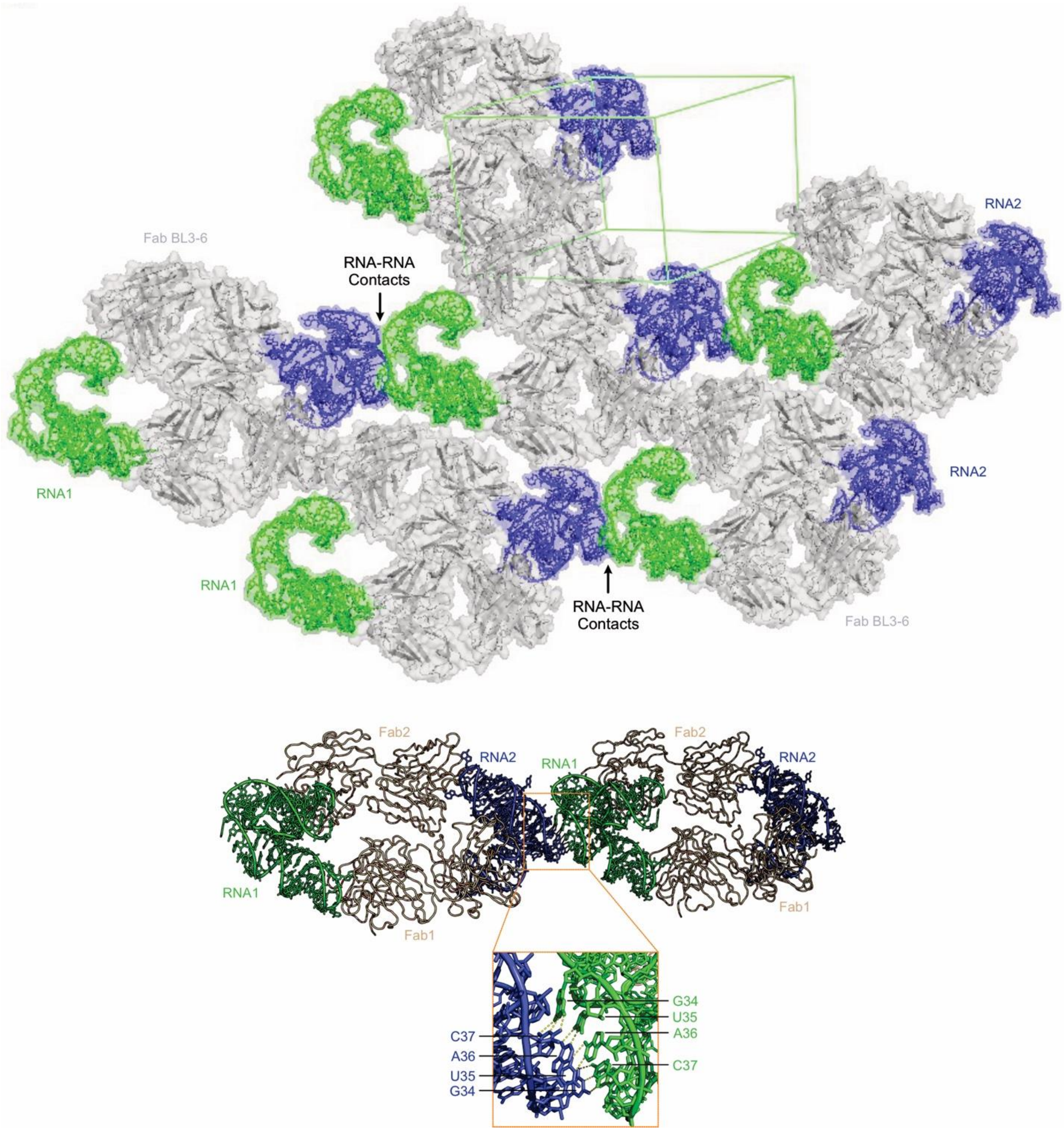


**Figure S6**. Crystal packing of the Fab-SLIIc complex showing the Fab- and RNA-mediated crystal contacts. Within the crystal lattice, including the Fab-RNA binding interface, the Fab-mediated contacts account for the majority of the crystal contacts, suggesting a critical role of the Fab in SLIIc crystallization. While there are no RNA-RNA contacts between the molecules within the asymmetric unit, the nucleotides G34 to C37 for each RNA molecule are found to be involved in base-pairing interactions with the symmetry-related molecule. The symmetry-related RNA molecules are labeled RNA1 (blue) and RNA2 (green) for clarity.

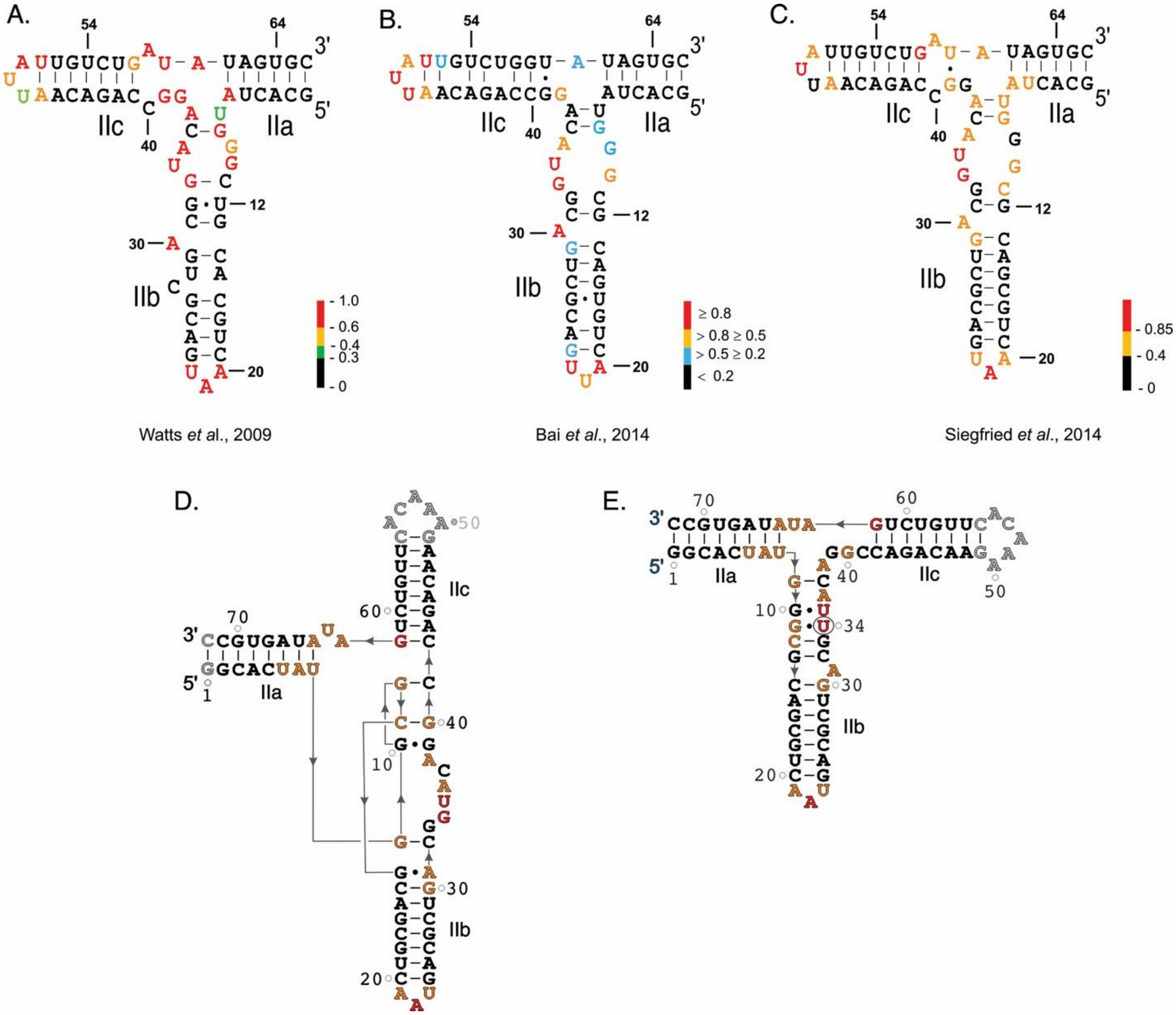


**Figure S7**. Predicted secondary structural model of RRE SLII showing the SHAPE reactivity profiles according to previous studies by (A) Watts *et al*. [1] in the context of the full-length HIV-1 genome, (B) Bai *et al*. [2] in the context of the minimal RRE construct (354 nts), and (C) Siegfried *et al*. [3] in the context of the entire HIV-1 genome (NL4-3 strain, ~ 9200 nucleotides). While the SHAPE reactivities of the IIb and IIc loop nucleotides are similar in all three studies, some variability is apparent for the nucleotides within or around the 3WJ, indicating the conformational heterogeneity of the junction. The representation of SHAPE reactivity profiles observed by Siegfried *et al*. within the crystal-derived secondary structure of the (D) SLIIc and (E) SLIIcG34U. These SHAPE reactivities are generally consistent with a mixture of these conformations in the solution, which is consistent with our Rev binding studies.

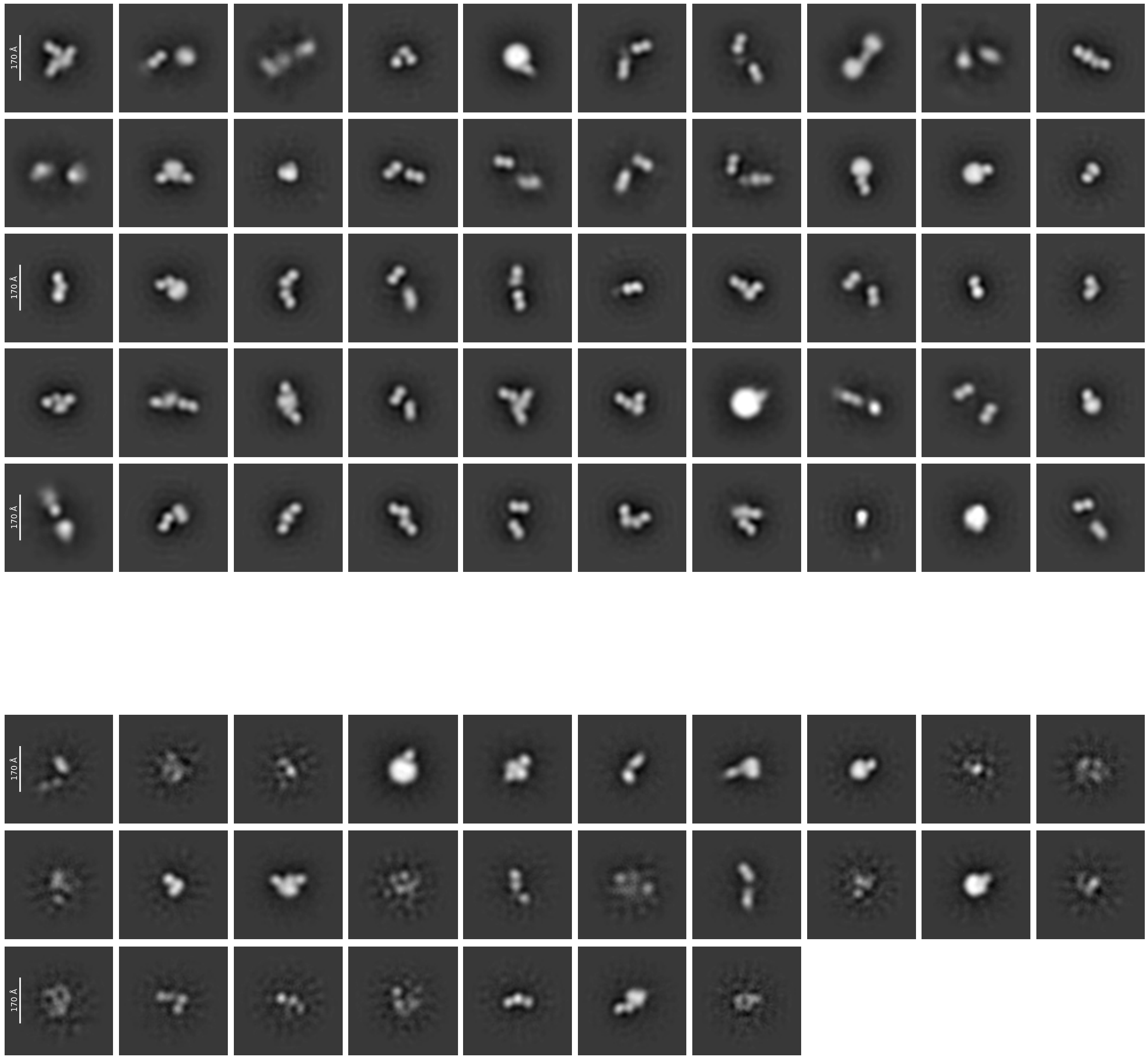


**Figure S8**. Negatively-stained electron microscopy (EM) images of the Fab-SLIIc complex particles at a 1.2 μg/ml concentration. Fifty 2D class averages for this concentrated sample show heterogeneous populations of monomers, dimers, and trimers of the Fab-SLIIc complex (top panel). Twenty-seven 2D class averages (bottom panel) were excluded due to the low quality of the particle images.

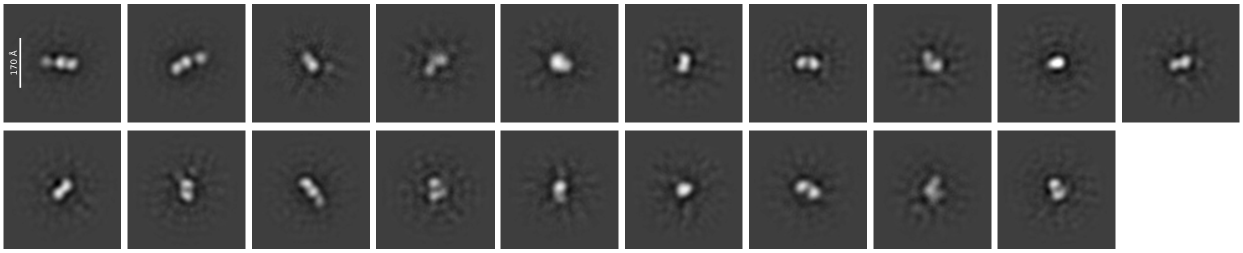


**Figure S9**. Negatively-stained electron microscopy (EM) images of the Fab-SLIIc complex particles at a 0.24 µg/ml concentration. Nineteen 2D class averages for this diluted sample show predominantly monomeric Fab-SLIIc complex particles.

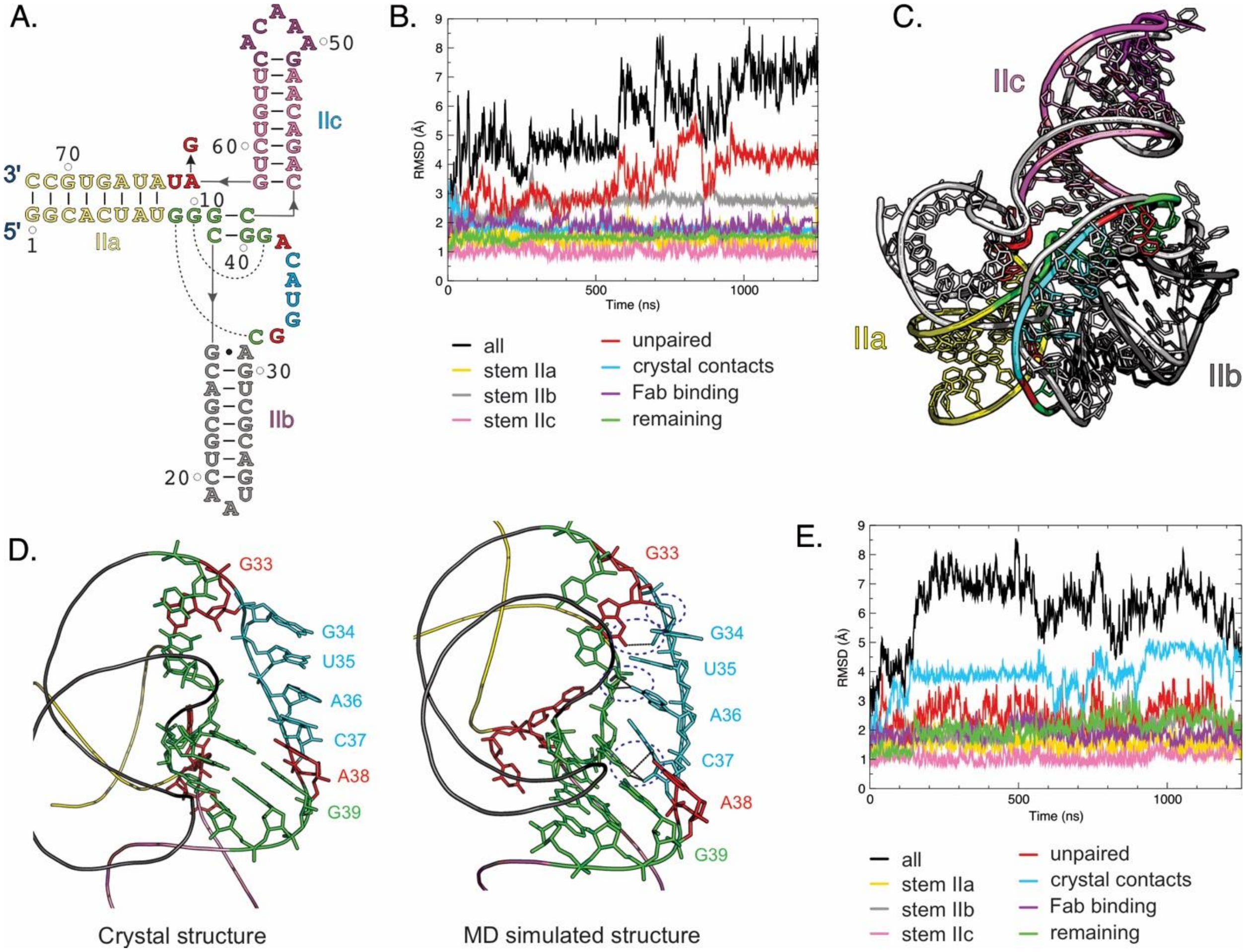


**Figure S10**. Molecular dynamics simulation analysis of the SLIIc crystal structure. (A) The crystal-derived secondary structure for the MD simulation. The stems IIa-c are colored yellow, gray, and pink, respectively. The unpaired nucleotides are colored red, and the nucleotides taking part in crystal contact are colored cyan. The nucleotides that are sequentially not part of the stem but structurally stacked on the stems are colored green, and the nucleotides involved in the Fab-binding are colored purple. The dotted curves represent long-range base-pairing interactions. (B) The root mean square deviations (RMSDs) of different structural components compared to the starting structure. Along the 1.25 µs (1250 ns) trajectory, frames were collected every 10 picoseconds, and RMSDs are plotted as running averages over 100 frames (i.e., 1 ns). (C) The superposition of the final simulated structure (colored according to the secondary structure) with the crystal structure (gray). (D) The structure of the crystal contact forming nucleotides (G34-C37) in the crystal (left) and after the simulation (right). The dotted blue ellipses highlight the nucleotides that create new hydrogen bonds (indicated by dotted lines). (E) The root mean square deviations (RMSDs) of different structural components compared to the starting structure generated based on a previous NMR model (see below Figure S11 for this NMR secondary structural model). All figure panels and corresponding labels, if any, are colored analogously for facile comparisons.

**Note**: The overall RMSD values for the system (Figure S10A) reach 8 Å and show significant variations, whereas the RMSD values of the three stems are small, with approximately 1.5 Å, 3 Å and 1 Å for stem IIa, IIb, and IIc, respectively (Figure S10B). This indicates that the source of high overall RMSD values is not the changes occurring at the stems. Similarly, the nucleotides that form crystal contacts display converged RMSD values overlapping in magnitude with the stem IIa. However, the RMSD for unpaired nucleotides (i.e., G33, A38, A63, U64) exhibit high fluctuations correlating with the overall RMSD values. These results suggest that the internal structure of the stems, crystal contacts, and the Fab binding domain are stable and remain close to the crystal structure. On the other hand, the unpaired nucleotides and the relative position of the stems undergo more significant dynamical fluctuations in solution that deviate from their relative orientations in the crystal structure (Figure S10C, RMSD of superposition = 10.2 Å). All the base pairs depicted in the crystal-derived secondary structure (Figure S10A) remained intact throughout the simulations. Interestingly, the four crystal-contact-forming nucleotides (G34-C37) in the simulated structure turned towards the RNA and formed non-canonical hydrogen bonds with nearby nucleotides – specifically, C37:O2 with A38:N6, A36:N6 with O2′ of G9 and C14, G34:N7 with G9:N2 (Figure S10D).

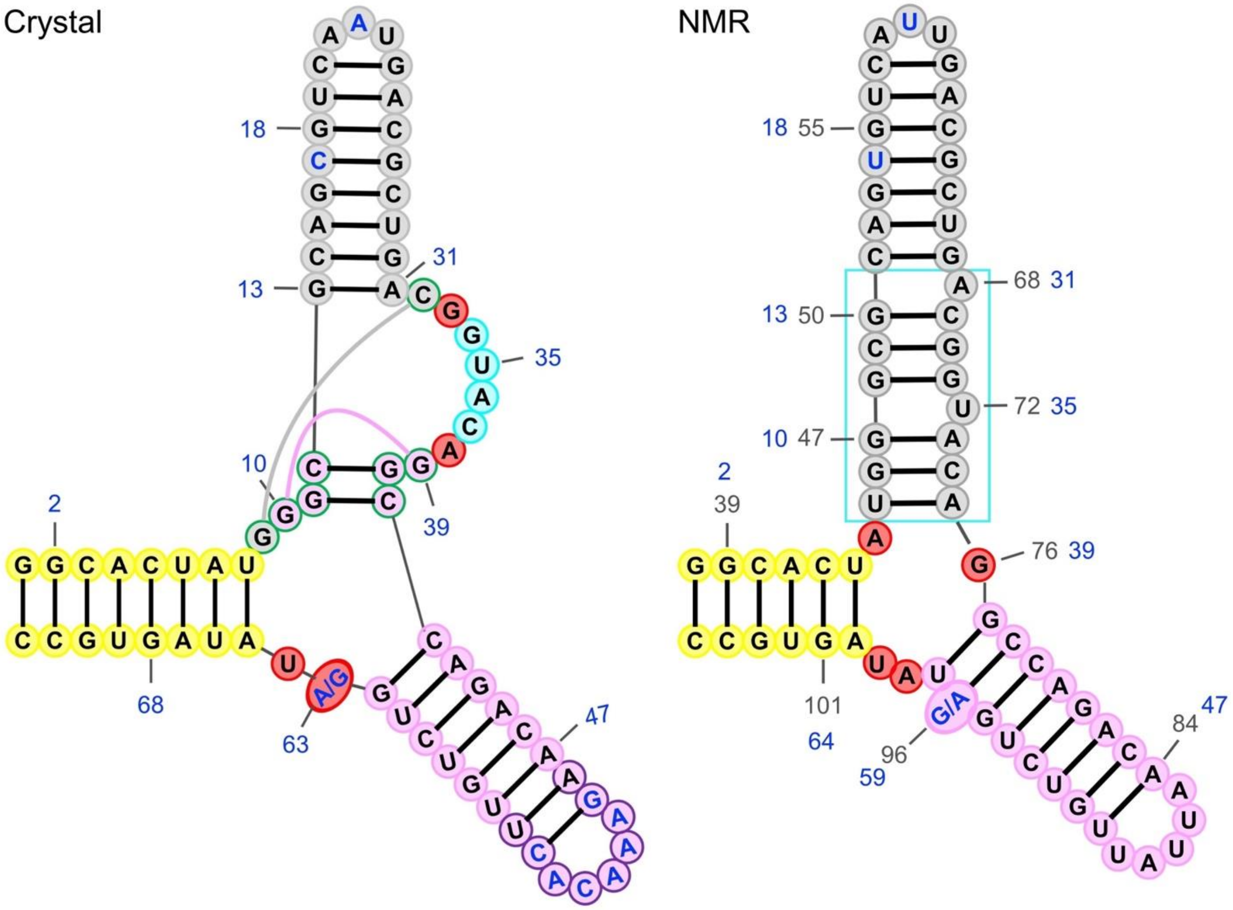


**Figure S11**. The secondary structure models of the SLIIc crystal structure and that proposed by the previous NMR study [4]. The stems IIa, IIb and IIc are yellow, gray, and pink, respectively; unpaired nucleotides are red, and the nucleotides in crystal contact are colored cyan. The nucleotides that are sequentially not part of the stem but structurally stacked on the stems are colored in the color of the corresponding stem but are highlighted with green outlines. The nucleotides corresponding to the Fab binding domain are outlined in purple. Nucleotides that differ between the two sequences are shown in blue. Canonical and sequential numberings are shown in gray and blue, respectively. To accommodate the length differences between the two sequences arising from the Fab binding sequence, numbers corresponding to the same nucleotide are displayed in both structures. These secondary structural models were prepared using the RNA canvas [5].

**Note**: One of the unpaired nucleotides, A63, appears as guanine in the previously proposed NMR model [4], which was predicted to form a Watson-Crick base pair with residue C41 (see Figure S11 for secondary structure comparisons). Given that A63 would not favor a Watson-Crick base pair with C, this could explain why the crystal structure base pairs and stems deviate from those suggested by previous NMR studies [4] with the IIb constructs. With this in mind, we modeled a mutation of A63 to G63 in the crystal structure and studied its dynamic behavior in solution. The resulting RMSD values for the A63G mutant were similar for the stems, with stem IIb converging to a slightly smaller value (Figure S10E). The unpaired group's RMSD values are lower but exhibit higher fluctuations. This is partly due to G63 forming a stable stacking interaction with the stem IIa without forming a base pair. Another difference regards the residues forming crystal contacts (G34-C37) that do not create as many new hydrogen bonds to hold them in place; however, this does not seem to be related to the mutation. All the base pairs depicted in the secondary structure map stayed intact for this A63G mutant simulation, similar to the simulations of the crystal structure. These results suggest that the intricate crystal structure has stable interactions and does not unfold or rearrange significantly in solution unless perturbated by mutation or Rev binding. Nevertheless, the several unpaired nucleotides within the 3WJ offer potential rearrangements of the base-pairing interactions when mutated, affecting the dynamicity of the region (RMSD values) and leading to a conformational change.

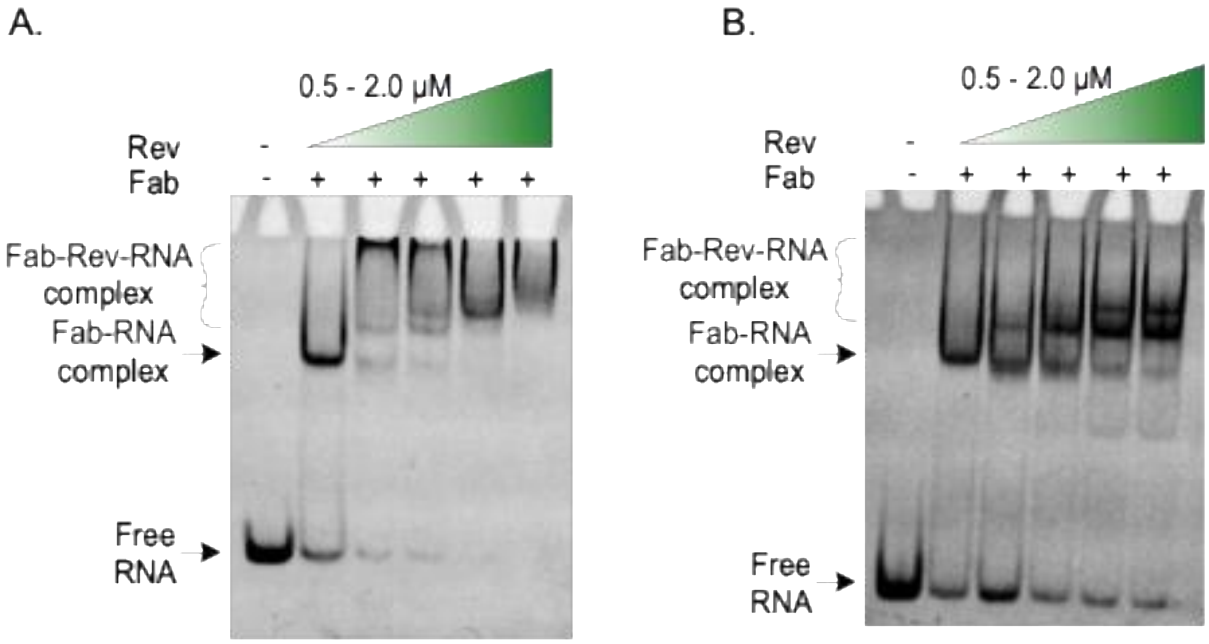


**Figure S12**. The native PAGE shows the Rev binding with the (A) SLIIcG34U and (B) SLIIcA36U crystallization constructs in the presence of the Fab BL3-6. Each lane was loaded with ~ 150 ng (~ 0.5 µM) of RNA and 1.5 µM of the Fab. The Rev protein concentration was varied from 0.5 – 2 µM as indicated by the gradient-filled green triangles. Both constructs yielded crystals, but SLIIcG34U-Fab complex crystals diffracted to 3.0 Å resolution.

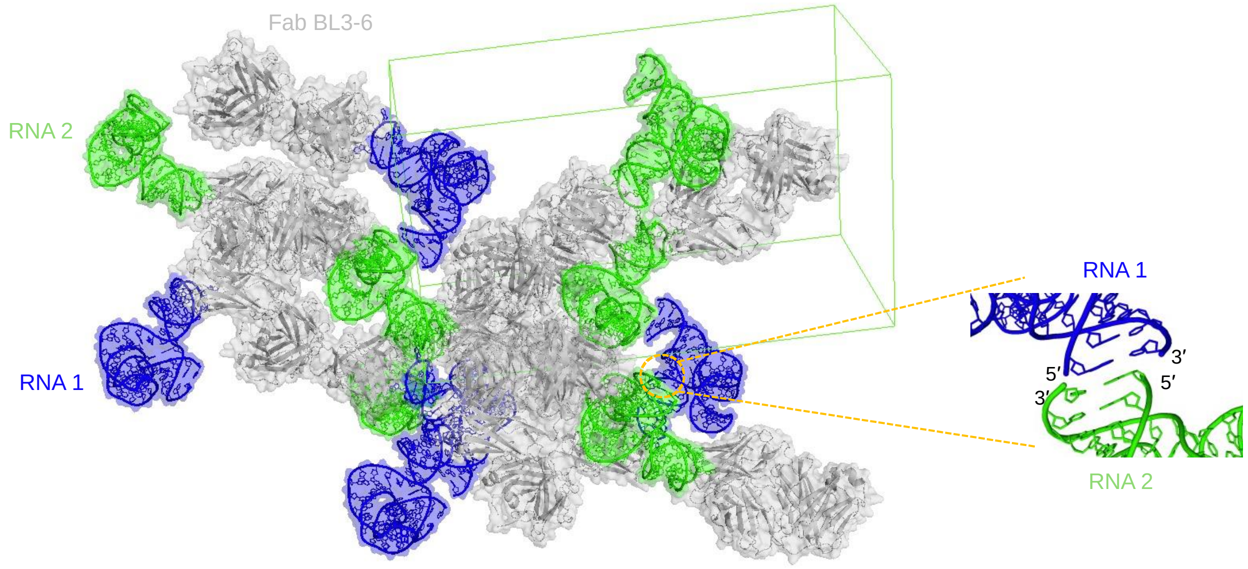


**Figure S13**. Crystal packing of the Fab-SLIIcG34U complex showing the Fab- and RNA-mediated crystal contacts. Within the crystal lattice, including the Fab-RNA binding interface, the Fab-mediated contacts account for the majority of the crystal contacts, suggesting a critical role of the Fab in SLIIc crystallization. However, the crystallographic asymmetric unit had a single Fab-RNA complex molecule, and the RNA-RNA crystal contacts involved the end-to-end helical stacking between the neighboring RNA molecules. The symmetry-related RNA molecules are labeled RNA1 (blue) and RNA2 (green) for clarity.

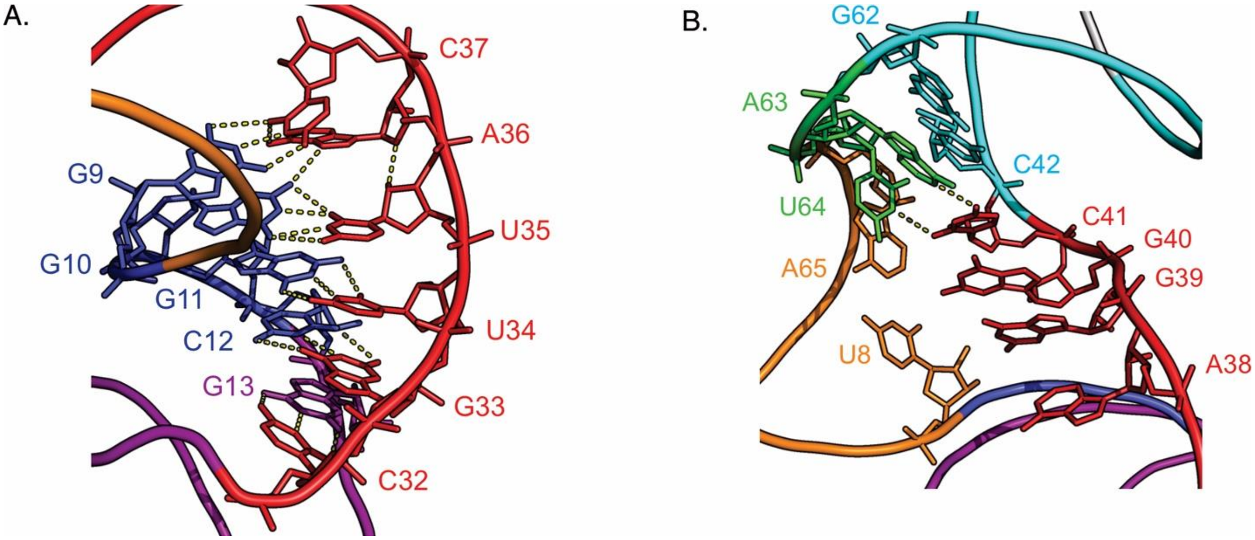


**Figure S14**. Specific interactions of nucleotides within the 3WJ of SLIIcG34U crystal structures. The Interactions of Jab G9-C13 nucleotides with (A) the Jbc G32-C37 and (B) Jca A63 and U64 nucleotides with the Jbc C41. The Jbc A38-C41 essentially remains unpaired within this SLIIcG3U 3WJ junction. Yellow dashed lines indicate heteroatoms within the hydrogen bonding distances.

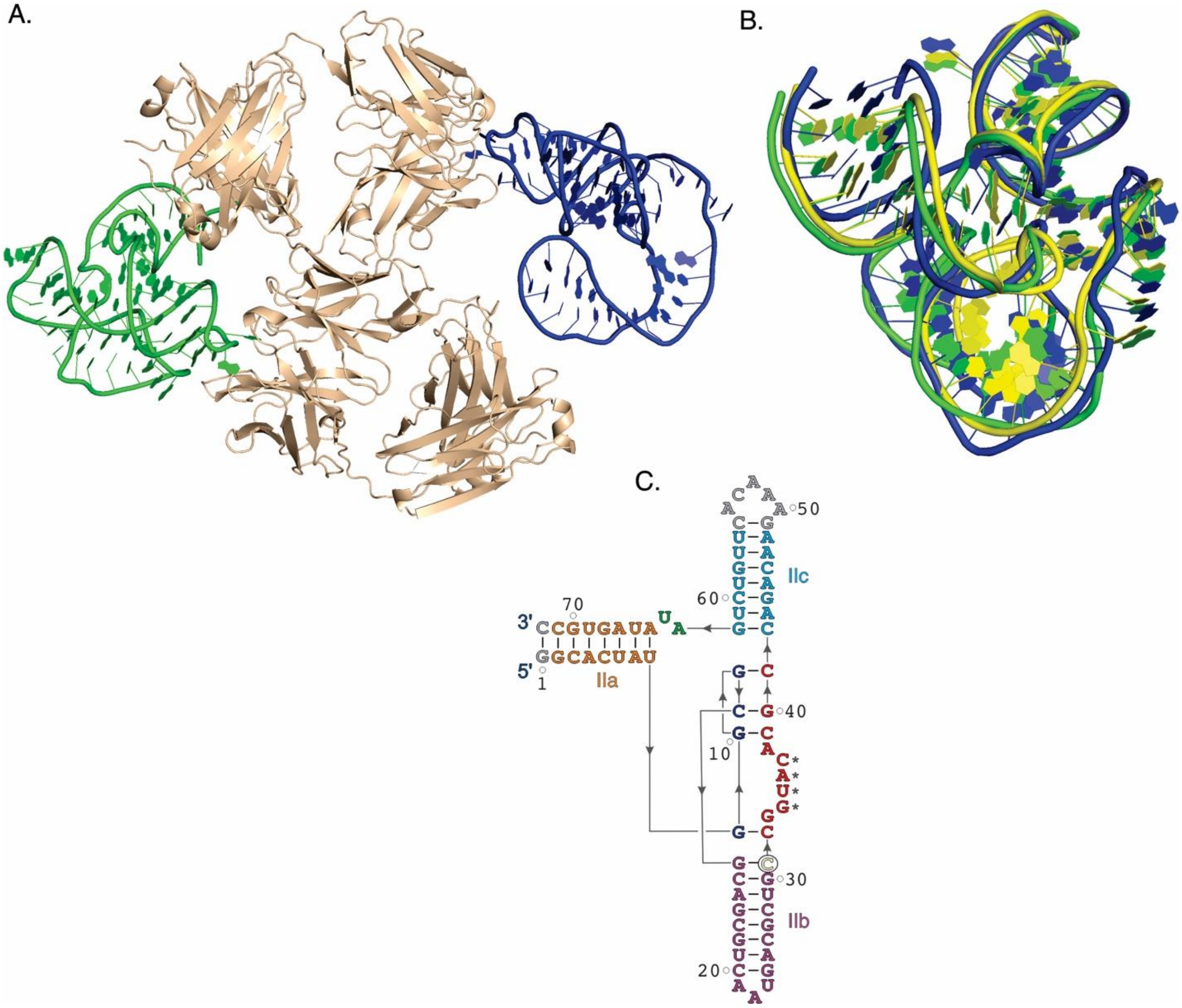


**Figure S15**. Crystal structure of the SLIIcA31C mutant in complex with Fab. (A) The crystallographic asymmetric unit contains two complexes similar to that in non-mutant SLIIc crystals. (B) superposition of the two RNA molecules (blue and green) identified within the crystallographic asymmetric unit (RMSD = 2.941 Å). This mutant structure is almost identical to the non-mutant (yellow) SLIIc structure (RMSD for superposition of SLIIc and SLIIcA31C = 2.963 Å). (C) The crystal-derived secondary structure of the SLIIcA31C shows the mutation position (colored yellow and circled) for facile comparison with the non-mutant SLIIc structure (Figure 1D).

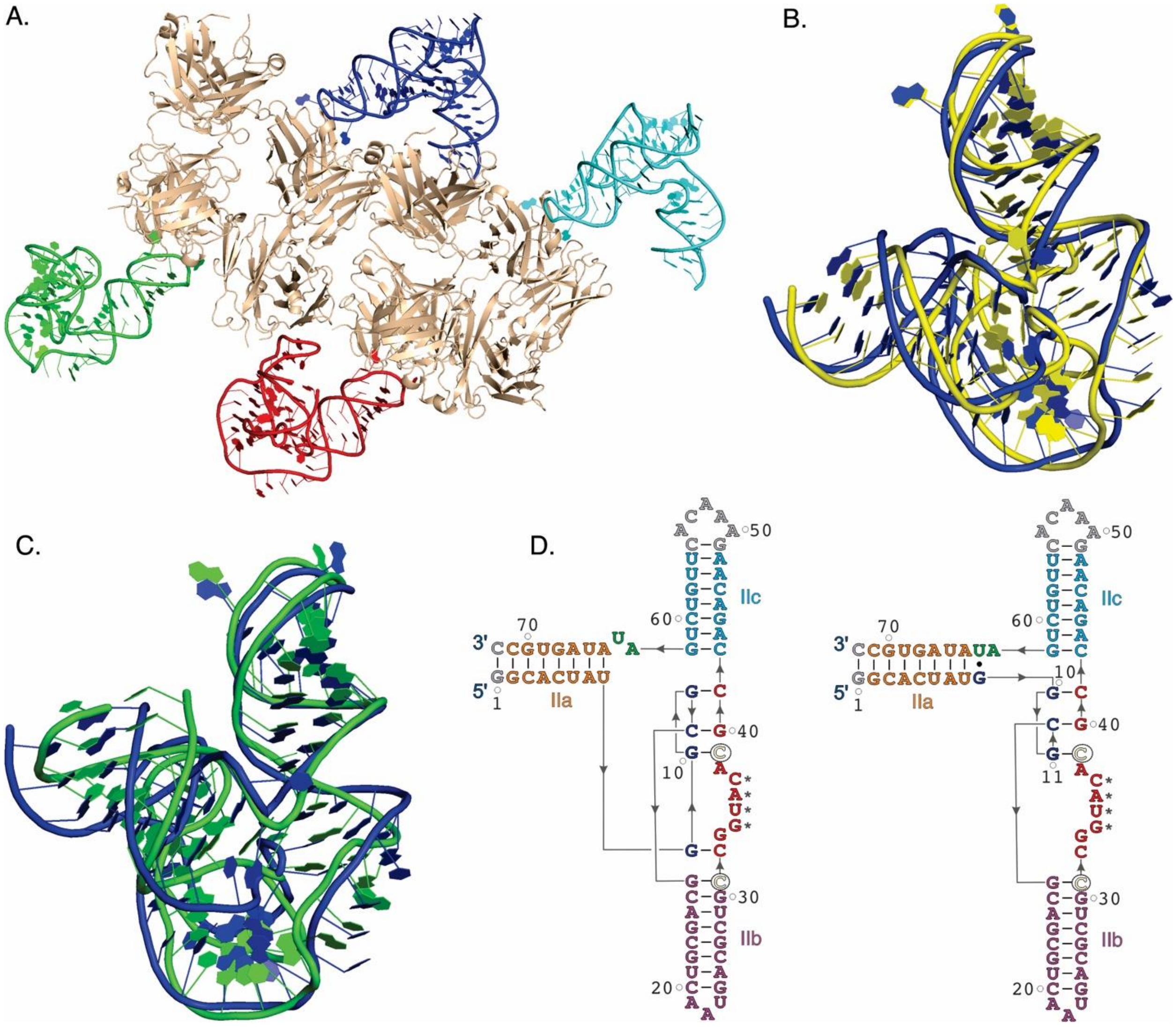


**Figure S16**. Crystal structures of the SLIIcA31C, G39C mutant in complex with Fab. (A) Unlike non-mutant SLIIc crystals, the crystallographic asymmetric unit contains four complexes. (B) Three of four SLII molecules (colored red, blue and cyan) adopted a similar conformation as the non-mutant SLIIc structure (RMSD for superposition of SLIIc, blue and SLIIcA31C and G39C mutant, yellow = 2.616 Å). (C) In contrast, one of the SLII molecules (colored green) has a slightly different configuration within the 3WJ (RMSD for superposition between these two structures, blue and green, within the asymmetric unit = 4.908 Å). (D) The crystal-derived secondary structures of the SLIIc A31C and G39C double mutant construct show the mutation positions (colored yellow and circled) for facile comparisons with the non-mutant SLIIc structure (Figure 1D). The secondary structures of both SLII conformations observed within the asymmetric unit are shown.

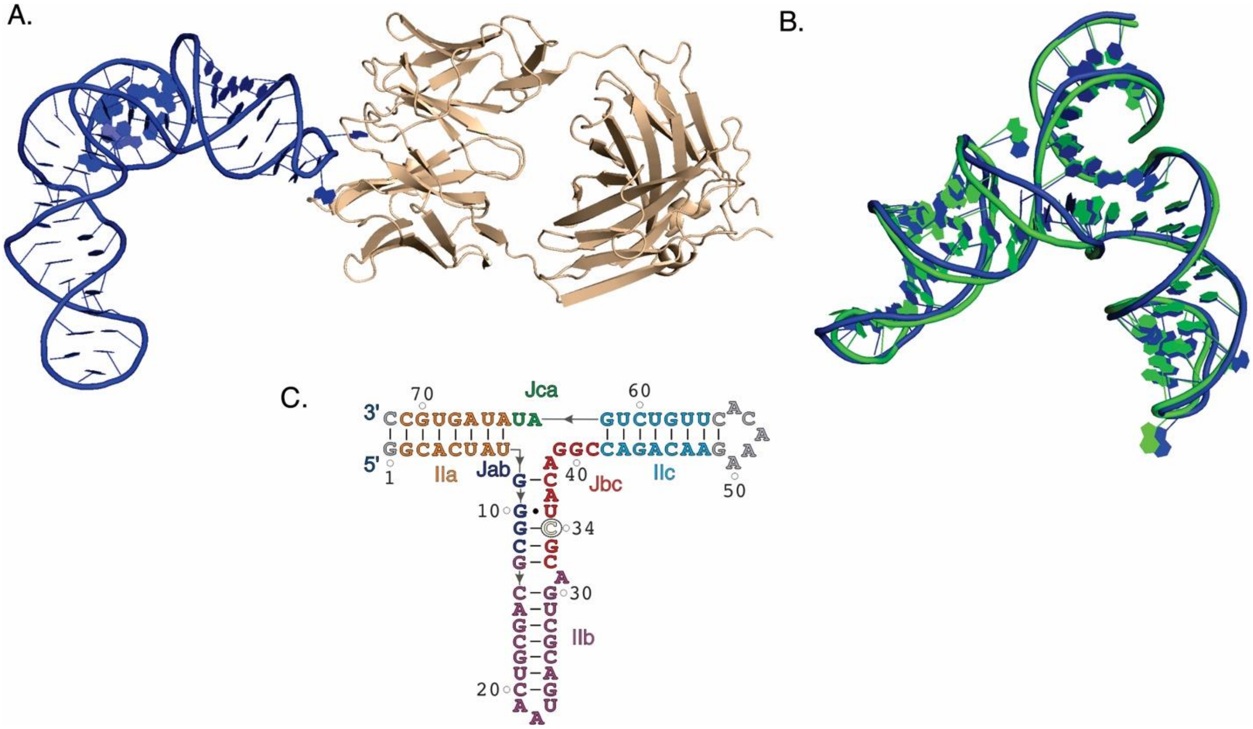


**Figure S17**. Crystal structure of the SLIIcG34C mutant in complex with Fab. (A) The crystallographic asymmetric unit contains a single Fab-RNA complex similar to that for the SLIIcG34U construct. (B) superposition of the two RNA structures for SLIIG34C (blue) and SLIIcG34U (green) (RMSD = 1.351 Å). (C) The crystal-derived secondary structure of the SLIIcG34C shows the mutation position (colored yellow and circled) for facile comparison with the non-mutant SLIIc (Figure 1D) and SLIIcG34U (Figure 4C) structures.

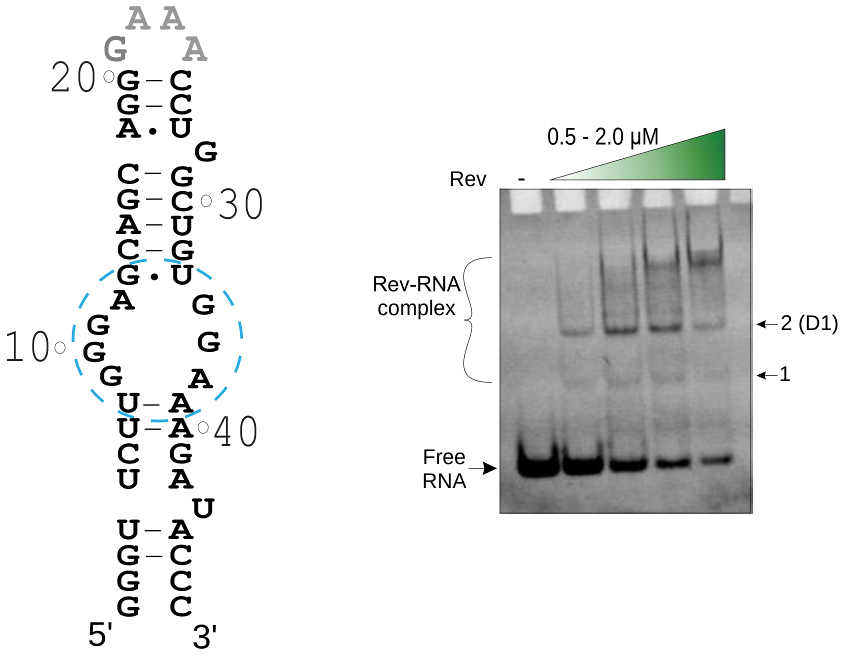


**Figure S18**. The native PAGE shows the Rev binding with an HIV-1 RRE stem-loop I construct. A GNRA-type tetraloop (gray) was used to cap the stem. The bulge within the stem (indicated by a dotted blue circle) has been shown to interact with Rev previously [6]. Our gel results suggest that stem-loop I will likely bind two Rev molecules (dimer) within the RRE structure. Each lane was loaded with ~ 150 ng (~ 0.5 µM) of RNA. The Rev protein concentration was varied from 0.5 – 2 µM as indicated by the gradient-filled green triangles.

### Supplementary Tables

**Table S1**. X-ray crystallographic data collection and structural model refinement statistics. The values in the parentheses are for the highest-resolution shell.

| ***Data collection*** | ***Constructs*** | | | | |
|---|---|---|---|---|---|
| | SLIIc (PDB: 9C2K) | SLIIc-G34U (PDB: 9C75) | SLIIc-G34C (PDB: 9E7G) | SLIIc-A31C (PDB: 9E7D) | SLIIc-A31C, G39C (PDB: 9E7E) |
| Space group | P1 | I222 | I222 | P1 | P1 |
| Resolution (Å) | 45.33-2.42 (2.45 – 2.42) | 34.64-3.04 (3.10-3.04) | 38.77-3.00 (3.05-3.00) | 34.2-3.10 (3.27-3.11) | 49.66-2.75 2.75 (2.75) |
| Cell dimensions a, b, c (Å) α, β, γ (°) | 72.36, 76.24, 82.60 116.56, 94.91, 102.75 | 90.88, 105.01, 186.60 90.0, 90.0, 90.0 | 90.78, 106.63, 187.33 90.0, 90.0, 90.0 | 72.87, 75.87, 83.51 63.17, 71.85, 77.75 | 84.37, 95.64, 111.24 77.48, 76.34, 74.25 |
| $R_{merge}$ (%) | 8.4 (0.392) | 18.3 (2.518) | 11.5 (1.671) | 87.5 (1.446) | 10.5 (1.866) |
| I/σI | 9.4 (1) | 7.1 (1.17) | 14.6 (1.0) | 1.5 (0.6) | 18.8 (0.9) |
| $CC_{1/2}$ | 0.978 (0.753) | 0.997 (0.383) | 0.998 (0.795) | 0.779 (0.319) | 0.832 (0.305) |
| Completeness (%) | 98 (94.4) | 99.8 (99.69) | 99.4 (94.67) | 98.9 (98.6) | 96.6 (93.3) |
| Redundancy | 1.8 (1.8) | 6.8 (6.9) | 13.6 (14.0) | 3.5 (3.4) | 3.6 (3.8) |
| ***Refinement*** | | | | | |
| No. reflections | 56456 (2247) | 17488 (2867) | 18631 (690) | 20060 (2906) | 82113 (2919) |
| $R_{free}$ (%) | 26.5 (0.3403) | 25.2 (0.4814) | 23.4 (0.5754) | 32.8 (0.3669) | 29.3 (0.3879) |
| $R_{work}$ (%) | 20.5 (0.3154) | 20.5 (0.3863) | 19.0 (0.4643) | 24.5 (0.2987) | 25.0 (0.3793) |
| *R.M.S. deviations* | | | | | |
| Bond angles (°) | 1.20 | 1.23 | 1.16 | 1.37 | 1.36 |
| Bond length (Å) | 0.010 | 0.009 | 0.008 | 0.010 | 0.010 |
| Average B-factor, all atoms ($Å^2$) | 79.0 | 135.0 | 144.4 | 63.5 | 132.2 |
| *Ramachandran plot of protein residues* | | | | | |
| Preferred regions (%) | 97.14 | 91.53 | 91.30 | 96.80 | 95.19 |
| Allowed regions (%) | 2.86 | 8.47 | 8.70 | 3.20 | 4.81 |

**Table S2**. Sequences of the RRE SLII constructs used in this study. The mutation positions are highlighted in red in bold font, and the Fab binding sequence is underlined.

| RNA constructs | Sequences |
|---|---|
| SLII WT | 5'GGCACUAUGGGCGCAGCGUCAAUGACGCUGACGGUACAGGCCAGACAAUUAUUGUCUGAUAUAGUGCC |
| SLIIc | 5'GGCACUAUGGGCGCAGCGUCAAUGACGCUGACGGUACAGGCCAGACAAGAAACACUUGUCUGAUAUAGUGCC |
| SLIIb | 5'GGCACUAUGGGCGCAGCGAAACACGCUGACGGUACAGGCCAGACAAUUAUUGUCUGAUAUAGUGCC |
| SLIIcG11A | 5'GGCACUAUGG**A**CGCAGCGUCAAUGACGCUGACGGUACAGGCCAGACAAGAAACACUUGUCUGAUAUAGUGCC |
| SLIIcG40A | 5'GGCACUAUGGGCGCAGCGUCAAUGACGCUGACGGUACAG**A**CCAGACAAGAAACACUUGUCUGAUAUAGUGCC |
| SLIIcG11A, C41U | 5'GGCACUAUGG**A**CGCAGCGUCAAUGACGCUGACGGUACAGG**U**CAGACAAGAAACACUUGUCUGAUAUAGUGCC |
| SLIIcG40A, C12U | 5'GGCACUAUGGG**U**GCAGCGUCAAUGACGCUGACGGUACAG**A**CCAGACAAGAAACACUUGUCUGAUAUAGUGCC |
| SLIIcA63G | 5'GGCACUAUGGGCGCAGCGUCAAUGACGCUGACGGUACAGGCCAGACAAGAAACACUUGUCUG**G**UAUAGUGCC |
| SLIIcG34U | 5'GGCACUAUGGGCGCAGCGUCAAUGACGCUGACG**U**UACAGGCCAGACAAGAAACACUUGUCUGAUAUAGUGCC |
| SLIIcA36U | 5'GGCACUAUGGGCGCAGCGUCAAUGACGCUGACGGU**U**CAGGCCAGACAAGAAACACUUGUCUGAUAUAGUGCC |
| SLIIcA63U | 5'GGCACUAUGGGCGCAGCGUCAAUGACGCUGACGGUACAGGCCAGACAAGAAACACUUGUCUG**U**UAUAGUGCC |
| SLIIcG39C | 5'GGCACUAUGGGCGCAGCGUCAAUGACGCUGACGGUACA**C**GCCAGACAAGAAACACUUGUCUGAUAUAGUGCC |
| SLIIcA31C | 5'GGCACUAUGGGCGCAGCGUCAAUGACGCUG**C**CGGUACAGGCCAGACAAGAAACACUUGUCUGAUAUAGUGCC |
| SLIIcA31C, G39C | 5'GGCACUAUGGGCGCAGCGUCAAUGACGCUG**C**CGGUACA**C**GCCAGACAAGAAACACUUGUCUGAUAUAGUGCC |
| SLIIcU64A | 5'GGCACUAUGGGCGCAGCGUCAAUGACGCUGACGGUACAGGCCAGACAAGAAACACUUGUCUGA**A**AUAGUGCC |
| SLIIcG11A, G40A | 5'GGCACUAUGG**A**CGCAGCGUCAAUGACGCUGACGGUACAG**A**CCAGACAAGAAACACUUGUCUGAUAUAGUGCC |
| SLIIc G11C, G40A | 5'GGCACUAUGG**C**CGCAGCGUCAAUGACGCUGACGGUACAG**A**CCAGACAAGAAACACUUGUCUGAUAUAGUGCC |
| SLIIcG34A | 5'GGCACUAUGGGCGCAGCGUCAAUGACGCUGACG**A**UACAGGCCAGACAAGAAACACUUGUCUGAUAUAGUGCC |
| SLIIcU14 insertion | GGCACUAUGGGCG**U**CAGCGUCAAUGACGCUGACGGUACAGGCCAGACAAGAAACACUUGUCUGAUAUAGUGCC |
| SLIIcA31C, G39C, G34C | GGCACUAUGGGCGCAGCGUCAAUGACGCUG**C**CG**C**UACA**C**GCCAGACAAGAAACACUUGUCUGAUAUAGUGCC |
